# Individual risk attitudes arise from noise in neurocognitive magnitude representations

**DOI:** 10.1101/2022.08.22.504413

**Authors:** Miguel Barretto Garcia, Gilles de Hollander, Marcus Grueschow, Rafael Polania, Michael Woodford, Christian C. Ruff

## Abstract

Humans are generally risk averse: they prefer options with smaller certain outcomes over those with larger uncertain ones. This risk aversion is classically explained with a concave utility function, meaning that successive increases in monetary payoffs should increase subjective valuations by progressively smaller amounts. Here, we provide neural and behavioural evidence that risk aversion may also arise from a purely perceptual bias: The noisy logarithmic coding of numerical magnitudes can lead individuals to *underestimate* the size of larger monetary payoffs, leading to apparent risk aversion even when subjective valuation increases linearly with the estimated amount. A formal model of this process predicts that risk aversion should systematically increase when individuals represent numerical magnitudes more noisily. We confirmed this prediction by measuring both the mental and neural acuity of magnitude representations during a purely perceptual task and relating these measures to individual risk attitudes during separate financial decisions. Computational model fitting suggested that subjects based both types of choices on similar mental magnitude representations, with correlated precision across the separate perceptual and risky choices. Increased stimulus noise due to the presentation format of risky outcomes led to increased risk aversion, just as predicted by the model. The precision of the underlying neural magnitude representations was estimated with a numerical population receptive field model fitted to the fMRI data of the perceptual task. Subjects with more precise magnitude representations in parietal cortex indeed showed less variable behaviour and less risk-aversion in the separate financial choices. Our results highlight that individual patterns of economic behaviour may, at least partially, be determined by capacity limitations in perceptual processing rather than by processes that assign subjective values to monetary rewards.

## INTRODUCTION

Risk aversion is the tendency of human and non-human decision-makers to choose smaller certain options over larger risky ones (Fox et al., 2015; Holt and Laury, 2002; Rabin and Thaler, 2001). While the population on average is risk averse, there is considerable variability in the individual strength of this tendency, and some people display risk-neutral or even risk-seeking behaviour (Bruhin et al., 2010; von Gaudecker et al., 2011). Traditional economic theories account for risk aversion by the non-linear, concave shape of the utility function that maps monetary outcomes to subjective utility of wealth (Kahneman and Tversky, 1979; von Neumann and Morgenstern, 1944). Such accounts thus conceptualise individual differences in risk aversion as differences in how the brain assigns subjective values to objective monetary outcomes. However, such theories fail to capture two key phenomena in real-life decision-making under risk (Mosteller and Nogee, 1951; Rabin, 2000).

First, if utility functions are concave enough to account for risk aversion in laboratory choices involving very small amounts of money, then decision makers should be risk averse even for gambles with very large potential gains and only moderate losses (Rabin, 2000; Rabin and Thaler, 2001). Human subjects, however, do not behave according to these assumptions. Second, many existing utility-based theories fail to explain the stochasticity in risky choice: Empirical evidence consistently shows that choices vary across repetitions of the same choice options (Gai and Vause, 2004; Hertwig et al., 2019; Hey, 2005; Khaw et al., 2021; Loomes and Sugden, 1995; Mosteller and Nogee, 1951). Click or tap here to enter text.While this phenomenon can be incorporated in models by simply adding a random error term to the utility function (Stott, 2006; Wilcox, 2008), such an approach fails to explain mechanistically why this choice stochasticity arises and whether it reflects some fundamental properties of neural computations that may lead to systematic biases and irrationalities.

Despite these conceptual problems, dominant neurocomputational accounts of individual differences in risky choice behaviour have mainly focussed on identifying neural valuation processes that may correspond to the computations captured by the utility function. While consistent correlations have been found in various brain prefrontal and subcortical regions (Christopoulos et al., 2009; Gilaie-Dotan et al., 2014; Heilbronner and Hayden, 2013; Platt and Huettel, 2008; Roitman and Roitman, 2010), it is still unclear from these findings what properties of neural processing may give rise to the individual variation in how the brain assigns value. This is particularly unclear since these computations are often conceptualized as a final stage in the value construction process that draws heavily on information passed on from preceding sensory and cognitive processing (Padoa-Schioppa and Conen, 2017; Spitmaan et al., 2019).

Here we test an alternative theoretical framework that explains risk aversion and stochasticity in risky choice not by idiosyncratic valuation processes, but as consequences of capacity restrictions and biases in the initial perception of the choice options. The core assumption of this framework is that even if decision makers take risky choices rationally – and thus attempt to maximize the expected value of the payoffs from their choices - they do not have access to the objective information about the option payoffs but only to capacity-constrained internal representations of it (Khaw et al., 2021). That percepts of numerical magnitudes are noisy and subject to several biases-resembling those observed for lower-level sensory percepts - has been well established by decades of work in psychophysics. For examples, perceptual judgments are stochastic (i.e., vary across repetitions) when human decision makers need to quickly estimate or remember the numerical magnitudes of a set of stimuli (Dehaene and Marques, 2002; Izard and Dehaene, 2008; Nieder and Miller, 2003, 2004; Nieder et al., 2002; van Oeffelen and Vos, 1982). Moreover, humans tend to increasingly underestimate larger (numerical) magnitudes in purely perceptual tasks (Anobile et al., 2012; Indow and Ida, 1977; Kaufman et al., 1949; Kramer et al., 2011; Krueger, 1972, 1984; Petzschner et al., 2015). Related studies in perceptual neuroscience suggest that this noise and biases in magnitude perception may be a direct consequence of the noisy and logarithmic way in which numerical magnitudes are encoded by neurons in parietal cortex (Harvey and Dumoulin, 2017; Harvey et al., 2015; Lasne et al., 2019; Nieder and Dehaene, 2009; Nieder and Miller, 2003; Piazza et al., 2004). This offers the intriguing possibility that from a neurocognitive perspective, individual differences in financial decision making may originate from biased perception originating in properties of parietal magnitude processing, rather than from subjective valuation processes instantiated in prefrontal and subcortical brain areas.

Recent economic models of risky choice have started to adopt this perspective and have proposed that risk attitudes may arise from the imprecision in mental representations of magnitudes (Frydman and Jin, 2022; Khaw et al., 2021). These models assume, in line with the literature on perceptual judgments (Petzschner et al., 2015), that logarithmic coding of the payoff information and Bayesian integration with an individual’s prior beliefs (shaped by more frequent exposure to smaller magnitudes) leads to more variable and systematically underestimated percepts for larger magnitudes (Petzschner et al., 2015). This has the consequence that even a decision rule that is adapted to maximize expected payoffs can result in choices that show hallmark patterns of risk aversion, in a manner that depends systematically on the noisiness of assumed magnitude representations (Khaw et al., 2021).

This perceptual account of risk aversion naturally accounts for two key empirical phenomena that utility-based models fail to explain. First, it naturally follows that economic choice will be stochastic, given that they are based on noisy mental magnitude representations (Woodford, 2020). Second, the logarithmic compression of mental magnitude representations can explain why subjects are risk averse even for arbitrarily small gambles (Rabin, 2000; Rabin and Thaler, 2001), since diminishing sensitivity for different payoffs should simply be a function of the log-*ratio* of potential payoffs (i.e., the distance on a logarithmic scale), irrespective of overall magnitude (Woodford, 2020). Crucially, the perceptual account of risk aversion also makes the novel prediction that individual or situational differences in risk aversion should be negatively related to the precision of the mental magnitude representations employed by the decision maker (Petzschner et al., 2015; Pouget et al., 2013; Woodford, 2020).

While these theories thus formalize how apparent risk aversion may emerge from biased perception and the noise in magnitude representations, empirical support for these theories is limited. In particular, the existing studies have fitted their model to a single economic choice task and inspected the relation of the fitted parameters with the choice variability in the same dataset (Frydman and Jin, 2022; Khaw et al., 2021). No study to date has linked any of these behavioural measures of risk preferences to characteristics of the neural coding of magnitudes. It is therefore unclear whether an individual’s risk aversion can indeed be traced back, in a mechanistic sense, to the acuity with which her brain represents magnitude information, and whether the noisiness (or inversely precision) of these magnitude representations is a stable trait that can parsimoniously account for the way in which an individual takes both perceptual and financial choices.

Here, we provide this evidence, by measuring the precision of mental and neural magnitude representations in a purely perceptual magnitude task and testing how these neurocognitive measures of perceptual magnitude precision can account for individual risk-taking behaviour in different contexts with varying sensory noise. By using a single unifying model capturing principles of magnitude representations (Izard and Dehaene, 2008; Khaw et al., 2021; van Oeffelen and Vos, 1982), we can thus not only link risk aversion to estimates of behavioural precision of mental magnitude perception, but also to its neural substrate of numerical representations in the intraparietal sulcus (IPS), which we measure with functional magnetic resonance imaging (fMRI).

## RESULTS

### The experiment

To test whether numerical acuity is an individual neurocognitive trait that (a) generalizes across perceptual and economic tasks and (b) determines individual differences in risk aversion, we devised an experiment comprising of two sets of tasks. First, we presented a perceptual magnitude comparison task in the MRI scanner during which subjects (*N* = 64) had to indicate which of two-coin clouds was more numerous (**Fig 1a**). This task allowed us to obtain both behavioural (Izard and Dehaene, 2008; van Oeffelen and Vos, 1982) and neural measures (van Bergen et al., 2015; Harvey et al., 2013) of numerical acuity. Then, in a second set of experiments outside of the scanner, we presented two sets of risky choice tasks to measure individual differences in risk aversion (**Fig 1b-c**). In one half of the risky-choice trials, we presented payoff magnitudes as coin clouds (**Fig 1b**) and thus in the same presentation format as also employed for the perceptual task. In the other half of trials, we presented the payoffs symbolically using Arabic numerals (**Fig 1c**). This allowed us to test whether individual differences in numerical acuity and risk aversion generalise across non-symbolic and symbolic settings, and whether risk aversion decreases if stimulus discriminability is increased (i.e., from non-symbolic to symbolic presentation). This latter hypothesis follows from model predictions (see below) that if internal noise in magnitude representations is reduced, any perceptual bias giving rise to risk aversion should also decrease. Variations in magnitude were matched across both risky-choice tasks, allowing explicit comparisons of precision between presentation formats (see **Methods** for details). To formally model behavioural data, we used a variant of a noisy logarithmic coding (NLC)-model (Khaw et al., 2021) that builds on established models of numerical cognition (Dehaene, 2003; Dehaene and Marques, 2002; Izard and Dehaene, 2008; van Oeffelen and Vos, 1982) and could be fitted to all choice tasks.

**FIGURE 1.**
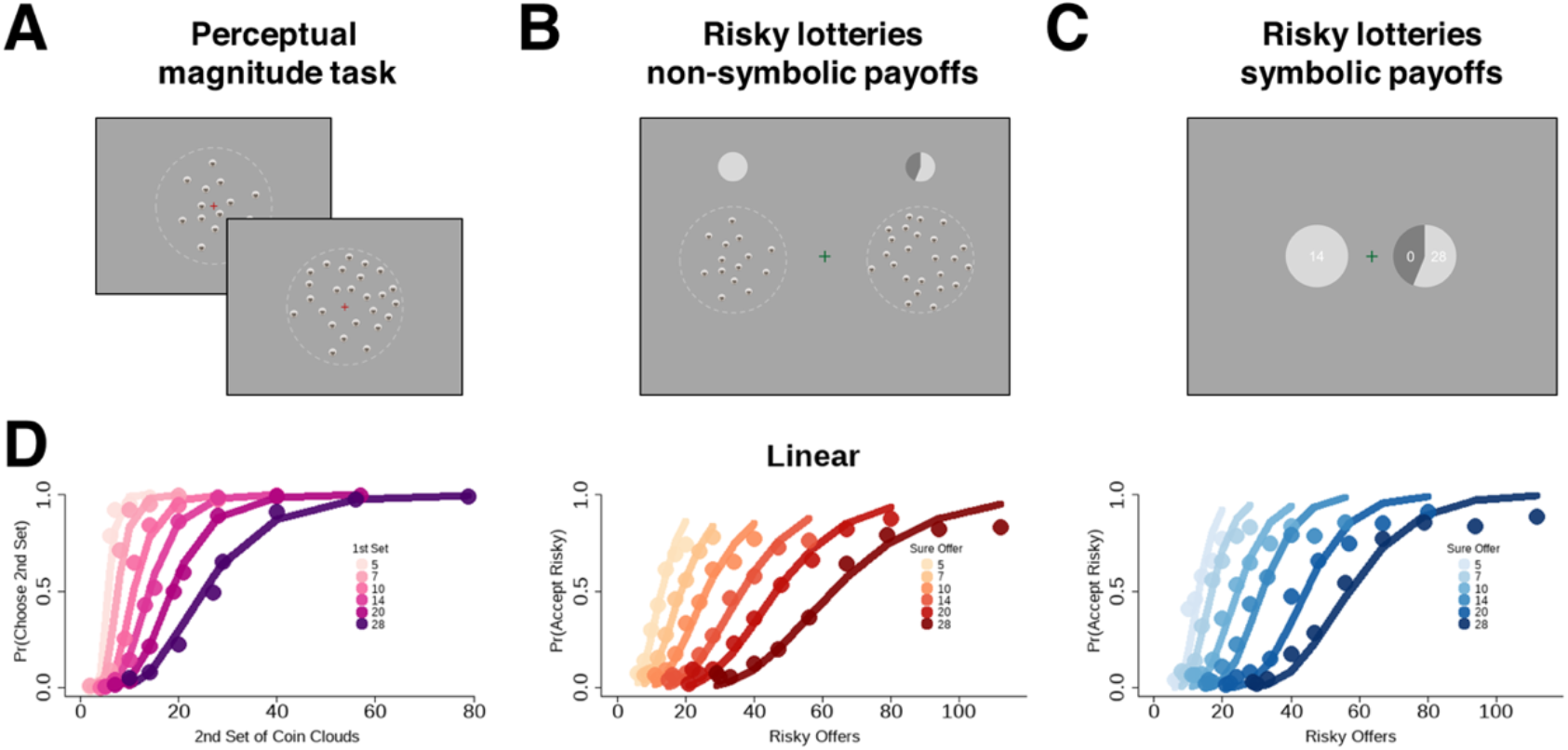
Experimental paradigm and behavioural results. **(a)** Participants performed a numerical decision-making task inside the MRI scanner. On every trial, two clouds of 1-CHF coins were presented sequentially, and participants had to indicate which of the two clouds contained more coins. We collected both neural and behavioural measurements to estimate the precision of the magnitude representations used for the task. Outside of the scanner, participants took risky choices in which they had to choose either a risky gamble or a sure offer. We visually displayed the potential payoffs of the offers as either **(b)** non-symbolic coin clouds similar to the perceptual task or **(c)** symbolic Arabic numerals. Probabilities were presented as pie charts, and we fixed the probabilities at *p* = 0.55 in favor of the risky monetary offer. **(d)** Observed probabilities of choosing the second stimulus for magnitude judgments (*left*) or choosing the risky option for risky choices (*middle* and *right*) plotted in linear space, separated by visual display type (non-symbolic payoffs, *middle* and symbolic payoffs, *right*). The six psychometric curves correspond to the magnitudes of the first stimulus arrays in the magnitude task or, analogously, the six possible sure offers in the risky-choice task.

### A common model for perceptual and risky choice

We employed the noisy logarithmic coding (NLC) model as the guiding framework to test the generalisability of numerical acuity across perceptual and risky choice tasks (see **Methods** for more details). The NLC posits that decisions maximize expected payoffs based on the mean Bayesian posterior magnitude estimate, which systematically integrates the prior belief about the magnitude distribution and the noisiness of the internal representations of the current magnitudes at stake (Petzschner et al., 2015). The model assumes that the brain represents the magnitudes of the risky payoff *X* and the certain payoff *C* by *r*_*x*_ and *r*_*y*_, two noisy estimates on a logarithmic internal scale modelled as samples from log-normal distributions,

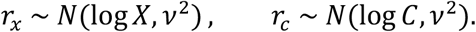

Perception is formalized as a Bayesian inference process that combines these internal estimates with a common prior, *X, C ~ N*(*µ, σ*^2^) that specifies the distribution of numerical magnitudes the decision maker expects to encounter in the testing environment. Central to the model is the individual parameter *ν* that represents the noise of the internal representations of numerical magnitudes. The parameter, *σ*, on the other hand, is the width of the prior that accounts for the dispersion of numerosities that the decision-maker deems plausible. The decision process involves optimization of the expected payoff based on the mean posterior magnitude estimate

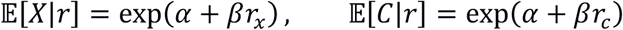

where *α* is a constant and 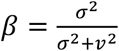 is the linear weighting of the prior versus the likelihood.

Here, the larger the noise in mental magnitude representations *r* (reflected by a larger *ν*), the shallower and more regressive the expected-value function over the payoffs.

With these underlying representational mechanisms, we can derive a psychometric function and predict the probability with which the decision-maker chooses *r*_*x*_ over *r*_*c*_,

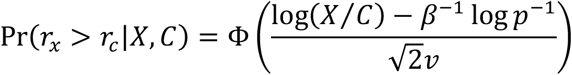

where Φ(·) is the standard normal cumulative density function and *p* is the probability of the risky payoff. In our task, *X* and *C* represent the objective magnitudes of the second- and first-coin clouds during the perceptual magnitude task and the magnitudes of the risky and certain payoffs during the risky choice task, respectively. We can conveniently estimate the NLC parameters using a standard probit model,

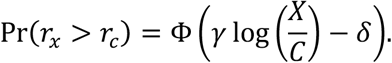

This parametrization is equivalent to the NLC via the relationships in the slope, 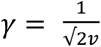, and intercept, 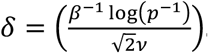.

The following are key measures for our analysis. First, the slope *γ* captures the precision in mental magnitude representations (the inverse of noise, *ν*) while the intercept *δ* captures the indifference point between both options. More precisely, the indifference point, 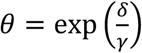, is where the individual is indifferent in choosing between *r*_*x*_ and *r*_*c*_. A crucial corollary of locating the indifference point is the ability to index risk aversion as risk-neutral probability, 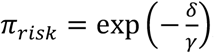, the probability level at which the decision maker should be indifferent between the risky and the safe option. The NLC prescribes that a decision-maker with imprecise mental magnitude representations should show *π*_*risk*_< 0.55 and thus behave as if the probability of receiving the risky payoff is smaller than it actually is (which, by definition, is apparent risk aversion). The “risk-neutral” probability during the perceptual magnitude comparison task, on the other hand, should trivially be *π*_*perceptual*_*≈* 1, since both options are associated with the same degree of (subjective) uncertainty.

### The NLC model is a better account for risky choice behaviour than established utility-based models

As a necessary condition, we established that our perceptual account of risk aversion can capture the empirical data at least as well as dominant economic theories. We used formal techniques to compare the NLC’s fit to the risky choice data with the fits of classical decision-making theories like expected utility maximization (EUM), cumulative prospect theory (CPT), and salience theory (ST), all in logit and probit form (see **Methods** for details). The formal model comparison revealed that the NLC fit the data best since it always had the lowest deviance information criterion (DIC), regardless of presentation format (see **Supplementary Fig 1a,b**). This underlines the plausibility of the neurocognitive operations formalized in the model.

### Decoding magnitude from neural activity

The NLC model assumes that both numerical magnitudes and potential payoffs are represented in the same noisy logarithmic manner. This builds on established neuroscientific findings that neural population activity in numerical parietal cortex (NPC) is tuned to numerical magnitudes, with the width of neural tuning reflecting moment-to-moment noisiness (or inversely, precision) of neural magnitude representations across repeated stimuli (Harvey et al., 2013; Piazza et al., 2004; Pouget et al., 2013). We now directly test these implicit, mechanistic links between neural and cognitive magnitude representations, by investigating the relationship between behavioural precision in the perceptual magnitude task and the fidelity of parietal magnitude representations measured by BOLD-fMRI during the perceptual decision-making task (van Bergen et al., 2015). To this end, we fitted a numerical population receptive field (nPRF) model that assumes that every patch of (parietal) cortex responds to a specific part of the number line, with its response profile characterized as a Gaussian kernel on the logarithmic number line (**Fig 2a**) (Dumoulin and Wandell, 2008; Harvey and Dumoulin, 2017; Harvey et al., 2013). We fitted the model in a cross-validated manner to a training set of 5 runs and then inverted it using Bayesian model inversion. This allowed us to obtain a posterior distribution over possible stimuli, given the BOLD activity patterns from the 6^th^, hold-out run (van Bergen et al., 2015). Thus, we could decode from the population response underlying the (previously unseen) BOLD activation patterns the most likely numerical magnitude of the presented stimulus (**Fig 2b**; see **Methods** for details).

**FIGURE 2.**
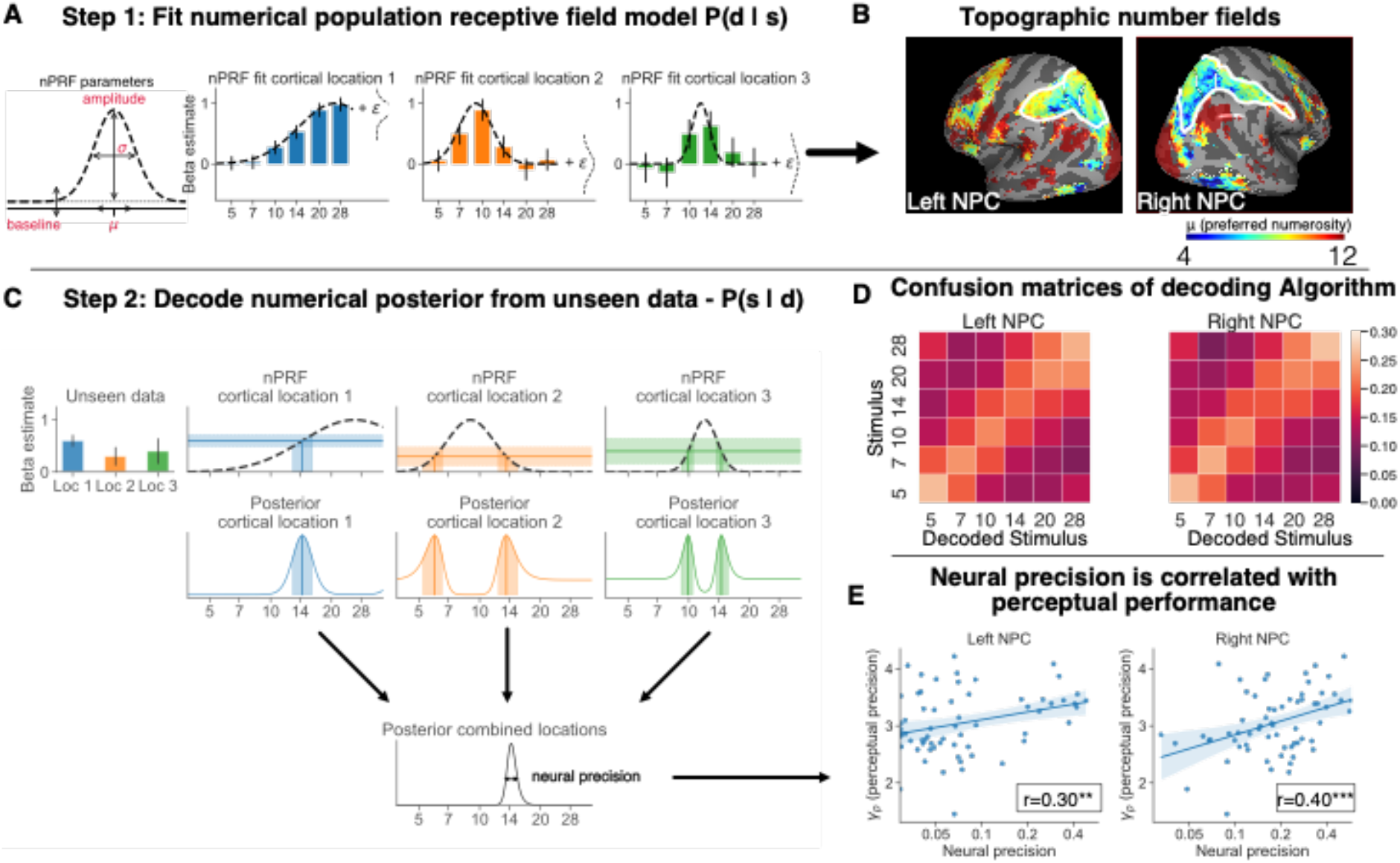
Decoding the precision of neural magnitude representations in parietal cortex. **(a)** The nPRF model fits a numerical, log-Gaussian receptive field (dashed curve lines) to BOLD activation patterns elicited by the magnitude of the first stimulus arrays in the perceptual task. In this hypothetical illustration, the first receptive field (RF, *blue*) activates stronger for larger magnitudes, the second RF (*orange*) strongly activates (roughly) between 7 and 10, and the third RF (*green*) between 10 and 14. **(b)** An illustration of decoding using Bayesian model inversion: The bar plots (*left, top panel*) are hypothetical beta estimates of BOLD activation responses to unseen stimuli. We invert the fitted nPRF model using Bayesian model inversion to obtain a posterior over possible numerical magnitudes of a stimulus, given a BOLD activity pattern. Our measure of imprecision in the neural magnitude representation is the standard deviation of this posterior (*middle panel*). The peaks of the posterior’s bimodal distribution (*solid vertical lines, middle panel*) represent the most likely stimulus, according to the BOLD activation in a single cortical location with a single receptive field (solid horizontal lines, top panels). The shaded areas correspond to the standard deviations of the beta estimates in the left panel. The posteriors of multiple cortical locations are then integrated into a single distribution that quantifies the posterior probability that different magnitudes have elicited the multivariate BOLD response pattern, as well as the uncertainty inherent in this assessment (bottom panel). **(c)** In line with earlier work, we found topographic organization of different numerosity preferences in bilateral numerical parietal cortex (NPC). **(d)** The confusion matrices of the numerical magnitude decoder show consistent and unbiased decoding of unseen numerosity stimuli. **(e)** The precision, 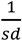, of the decoded posteriors is significantly correlated with the precision of mental magnitude representations employed during the perceptual decision-making task, as indexed by the parameter *γ*_*perceptual*_ * *p* < 0.05, ** *p* < 0.01, and *** *p* < 0.001.

In line with previous findings of topographic neural magnitude representations in parietal cortex (Harvey and Dumoulin, 2017; Harvey et al., 2013, 2015), we found that large regions in parietal cortex were sensitive to numerical magnitude stimuli, and that many of these were topographically organised according to their most-preferred numerosity (**Fig 2c**). Given established links between neural decoding accuracy in IPS and task performance (Kersey and Cantlon, 2017; Lasne et al., 2019), we manually segmented a region in IPS that was both magnitude-sensitive and showed topographic organisation, which we labelled left and right NPC. The neural magnitude representations in this NPC area were indeed highly specific, as we ascertained by inverting the encoding model and decoding the presented stimulus from NPC-BOLD activity patterns in held-out data. This worked better than chance (16.67%) in 56 out of 64 subjects in left NPC (average accuracy 23.6%, *s*.*d*. 5.4%) and 60 out of 64 subjects in right NPC (average accuracy 23.8%, *s*.*d*. 5.2%). These accuracies were significantly different from chance level decoding (16.67%) at the group level (*t*(63) = 10.4, *p* < 0.001 for left NPC and *t*(63) = 11.4, *p* < 0.001 for right NPC). Reassuringly, misclassified stimuli were usually classified as a stimulus relatively close in magnitude (**Fig 2d**). Moreover, in line with the notion that magnitude representations may be somewhat right-lateralized (Lasne et al., 2019), the correlation between the presented magnitude and the mean decoded posterior was higher for right NPC (*r* = 0.216) than for left NPC (*r* = 0.117) (paired *t*-test: *t*(63) = 6.17, *p* < 0.001), and the standard deviations of the decoded posterior were smaller (*right*: 6.9 vs *left*: 17.4). Overall, in line with our expectations, we could identify topographic areas in parietal cortex for each individual that reliably encoded numerical magnitudes in the perceptual task.

### Behavioural differences in numerical acuity relate to the precision of neural magnitude representations

Crucially, our neural model allowed us to not only identify neural magnitude representations in parietal cortex but also to derive a standard index of how precisely magnitudes are represented by neural population activity (see van Bergen and Jehee, 2019; van Bergen et al., 2015; Li et al., 2021; Walker et al., 2020): We can measure each individual’s general degree of *neural precision* in encoding magnitudes by the mean precision, 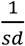, of the posterior across all stimulus categories (**Fig 3a**, *red arrow*). The NLC model implicitly assumes that its measure of precision in mental magnitude representations should reflect the precision of the corresponding neurobiological representations. Consistent with this hypothesis, we found significant positive correlations between each subject’s performance on the perceptual task, measured by the precision parameter, *γ*_*perceptual*_, and the index of general neural precision, 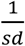 of decoded posteriors (averaged over all 6 stimuli categories) in both left and right NPC (*r*_*left*_ = 0.32, *r*_*right*_ = 0.42, both are *p* < 0.001; **Fig 2e**). This confirms a strong link between the noise in neural representations of numerical magnitudes and variability in performance during magnitude perception, particularly for right NPC (Kersey and Cantlon, 2017; Lasne et al., 2019).

**FIGURE 3.**
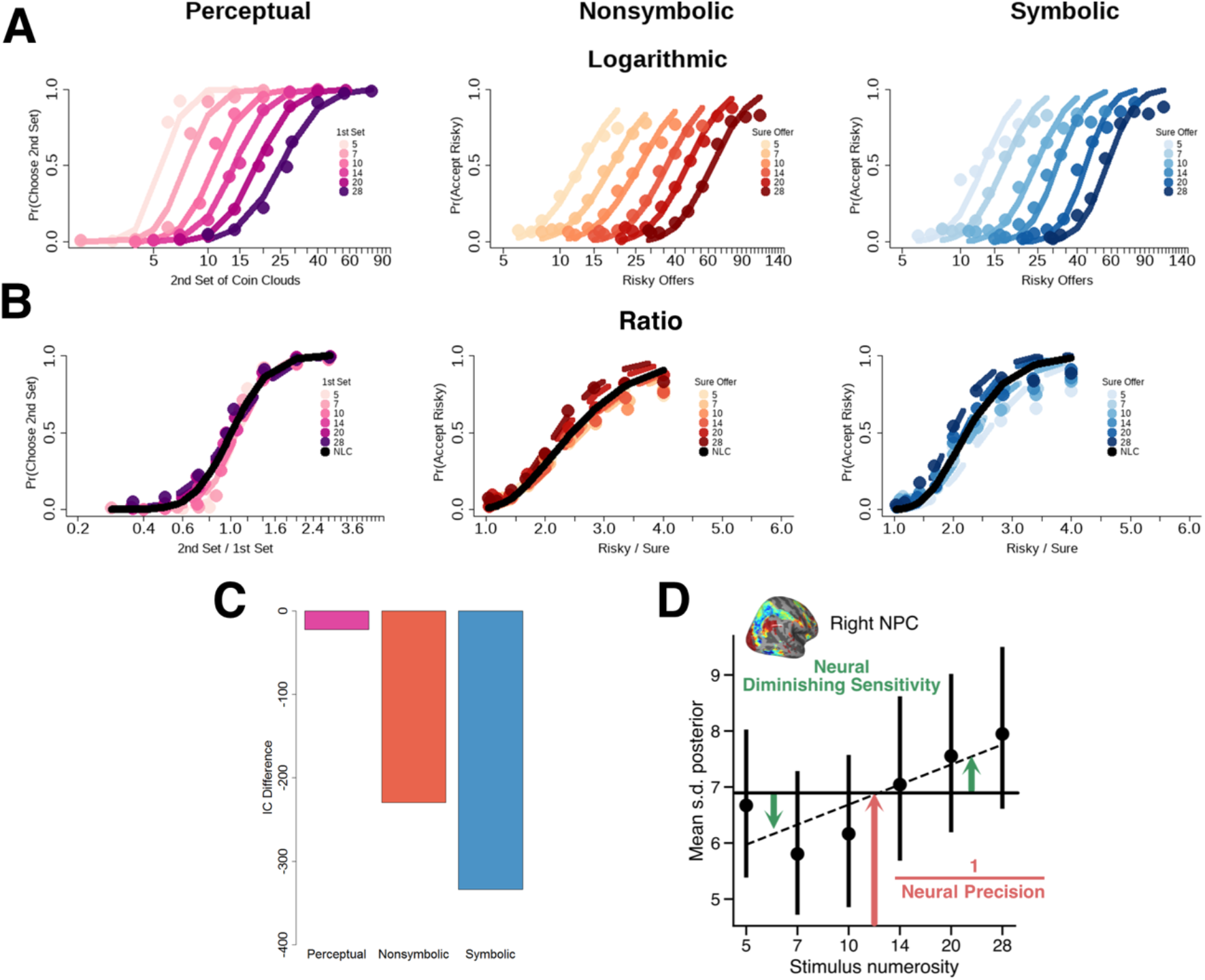
Domain-generality of scale invariance across perceptual and risky choice tasks. **(a)** The same observed choice probabilities as in Fig 1(d) but plotted in logarithmic space and **(b)** as the ratio of the second- and first-coin cloud magnitudes (magnitude task) or of the risky and sure payoffs (risky-choice task), to test for scale invariance; initial inspection indicates that the slopes of these psychometric curves are similar irrespective of task domain or visual display, already suggesting common logarithmic magnitude coding across tasks. Moreover, the slopes of the six psychometric curves stack over each other, and we can fit a single psychometric curve (*solid black curve*) to account for all choice probabilities in all task domains and visual displays. **(c)** Difference in deviance information criterion (DIC) between the best model (the NLC with one slope parameter for all experimental conditions) and the competing unrestricted model with separate slopes for each combination, for all types of choices and displays. **(d)** The dispersion (*s*.*d*.) of the decoded posteriors increases as a function of stimulus magnitude—a hallmark of scale invariance. We quantified, on a subject-to-subject basis, both the average *neural precision* (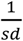 of the posterior) across magnitudes, as well as the n*eural diminishing sensitivity*, the extent to which neural representations get noisier as a function of stimulus magnitude.

### Evidence that similar logarithmic magnitude representations may guide both perceptual and risky choices

A central assumption of the NLC is that noisy logarithmic magnitude representations should be scale invariant, which should be evident in at least two ways: (a) the probability of choosing the second coin cloud in the perceptual task and the risky gamble in the risky choice task should be determined by the log-ratio of the two magnitudes, 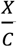, rather than by their absolute magnitudes; and (b) the noise in internal magnitude representations should increase with magnitude. To visually test for these hallmark signs of the representations assumed by NLC, we plotted the probabilities of judging the second set of coins larger than the reference stimulus (**Fig 1a**) or choosing the risky gamble over the sure gamble (**Fig 1b,c**) in both linear (**Fig 1d**) and log (**Fig 3a**) spaces. Initial visual inspection already shows that in linear space, the choice curves vary differently and are skewed. But when replotted on a log scale, the slopes (*γ*) become very similar and even more so in log-ratio scale, where all choice curves are practically identical (**Fig 3b**), suggesting scale-invariance. We also formally tested for scale invariance by comparing model fits of a single NLC psychometric curve to an unrestricted model that separately fitted six such psychometric curves for each of the six reference stimuli. Model comparisons revealed that the NLC’s single psychometric curve fitted to all magnitude stimuli explained the data far better than the unrestricted model (**Fig. 3c**), thus confirming scale invariance in the choice data.

Crucially, we also tested whether scale invariance applies to the neural magnitude representations identified with our encoding-/decoding approach. In addition to the precision of neural representations across all magnitude levels (mean precision, 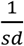, **Fig 3a**, *red arrow)*, we could also derive measure of *neural diminishing sensitivity (DM)*: The regression slope of the decoded posterior’s mean *sd* on stimulus magnitude, indexing how strongly, in a given individual, neural representations become less precise with increasing magnitudes (**Fig 3a**, *green arrows)*. Note that the first measure (*neural precision*) can be contaminated by non-cognitive factors unspecific to magnitude coding, such as size of the cortical sheet, neurovascular coupling, attention, and head movement (Naselaris and Kay, 2015; Snoek et al., 2019). The second measure (*neural DM*) is less likely to be affected by such unspecific general noise factors, which presumably do not vary systematically with trialwise stimulus magnitudes. Thus, for subjects who exhibit imprecision specifically in the neural coding of magnitudes, and for whom this coding follows scale invariance, we expect the slope of decoding uncertainty across magnitude levels to be systematically higher, on top of any general neural noise also affected by non-cognitive noise sources.

In line with these predictions, we found that the larger the stimulus magnitude, the noisier the neural magnitude representation (in natural space; repeated-measures correlation; for both left NPC (average *r* = 0.23, *p* = 0.001) and right NPC (**Fig 3d**, average *r* = 0.43, *p* < 0.001). This also confirms scale invariance in neural magnitude encoding and suggests that decisions in all tasks and contexts were guided by similar neurocognitive magnitude representations.

### The NLC captures how noise in magnitude representations mechanistically leads to perceptual bias and risk aversion

A crucial implication of the NLC model is that it specifies the link between magnitude representation noise and the point at which participants become indifferent between the choice options. In perceptual magnitude, where it is known a priori that there is no outcome uncertainty (i.e., probabilities are equal to 1), the individual’s indifference point should lie around *θ*_*perceptual*_ ≈ 1, the point where both magnitudes are equal, *X* = *C*. In risky choice, the objective outcome probability *p =* 0.55 should lead to an indifference point 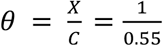 for perfectly precise magnitude representations (*ν* = 0). However, any degree of non-zero noise (*ν* > 0) will shift the indifference-point threshold to 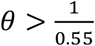 and thus to apparent risk aversion (since people become indifferent only for larger *X* at the given *p*). This prediction by the NLC should be evident by the location of the probit intercept, which should not be statistically different from zero, *δ* = 0, for perceptual judgements but greater than zero, *δ* > 0, for risky choice.

Indeed, population-level posterior distributions of the intercept estimated with a hierarchical Bayesian framework were all in line with model prescriptions. (**Supplementary Fig. 2a and 5a**, *δ*_*perceptual*_ = −0.168, *p*_*mcmc*_ = 0.204; **Supplementary Fig. 2b**, *δ*_*nonsymbolic*_ = 2.41 and *δ*_*symbolic*_ = 2.97, *p*_*mcmc*_< 0.001 for both measures). Post-hoc tests revealed that the indifference-point threshold for perceptual magnitude was no different from 1 (**Supplementary Fig. 2c**, *θ*_*perceptual*_ = 0.95, *p*_*mcmc*_ = 0.204) while the indifference points for risky payoffs across visual presentation formats were greater than 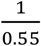 (**Supplementary Fig. 2d**, (*left panel*), *θ*_*nonsymbolic*_ = 2.46 and, *θ*_*symbolic*_ = 2.24, *p*_*mcmc*_< 0.001 for both measures). Given that previous work (Khaw et al., 2021) only tested the NLC within a single economic choice task, our results extend the NLC’s generalisability as a normative model that can flexibly account for biases in both perceptual and risky economic choices.

### Similar magnitude representations guide risky choice across visual display formats and individuals

The NLC defines the noisiness of mental magnitude representations *ν* as a function of *γ*, the precision with which individual decision-makers can mentally represent magnitude stimuli. At the same time, the NLC assumes that choice behavior should independently be influenced by different beliefs about the variability of magnitudes in the environment (measured by the width of the prior, *σ*) as well as by sensory noise in the stimulus display (which should also affect the precision of magnitude representations). However, these core features of the NLC remained thus far untested, since previous studies that have related risk aversion, *π*, to the precision in mental magnitude representation, *γ*, have done so with parameters fitted to a single economic task with only one type of visual display (e.g., symbolic payoffs). If *γ* really reflects an individual trait, then the relationship between *π* and *γ* should be robust to variations in sensory noise; but the sensory noise level should nevertheless have independent systematic effects on risk aversion.

We tested this hypothesis by comparing the same risky choices presented either symbolically as numbers or non-symbolically as coin clouds (**Fig 1b,c**). First, we predicted that individuals should be more risk-averse when faced with non-symbolic payoffs, since symbolic payoffs are easier to identify and map onto mental magnitude representations (note that by matching the distribution of stimulus magnitude of non-symbolic and symbolic payoffs, the prior between the two stimulus displays should be no different). Second, we expected that we will replicate previous results of a positive and nonlinear relationship between risky payoff precision and risk aversion (Khaw et al., 2021) for both display formats, but we crucially predicted that each individual’s magnitude noise will be related across both visual display formats.

Our results confirm all these predictions. Population-level posterior distributions of the corresponding parameters confirmed that representations of non-symbolic payoffs were less precise than symbolic ones (**Fig 4a**, *γ*_*nonsymbolic*_ = 2.76, *γ*_*symbolic*_ = 3.56, *p*_*mcmc*_< 0.001) and that the individual indifference points between risky and sure gambles were indeed higher during non-symbolic payoff presentations (*θ*_*nonsymbolic*_> *θ*_*symbolic*_ (**Supplementary Fig 2d** (*right panel*), *p*_*mcmc*_ = 0.003). As expected given the noisiness of mental magnitude representations, risk aversion was substantial for both display formats (**Fig. 4c** (*top-right panel*), *π*_*nonsymbolic*_ = 0.408, *π*_*symbolic*_ = 0.448 are less than 0.55, *p*_*mcmc*_< 0.001) and risk-neutral probabilities were indeed systematically smaller (corresponding to more risk aversion) for non-symbolic payoff display formats (**Fig. 4c** (*middle-right panel*), *p*_*mcmc*_ = 0.003). Importantly, these differences in the indifference point and risk-neutral probability did not appear to reflect different beliefs about stimulus magnitude distributions, since the estimated priors were not different between non-symbolic and symbolic payoffs (**Fig. 4b**, *σ*_*nonsymbolic*_ =0.366, *σ*_*symbolic*_ =0.343, *p*_*mcmc*_ = 0.204).

**FIGURE 4.**
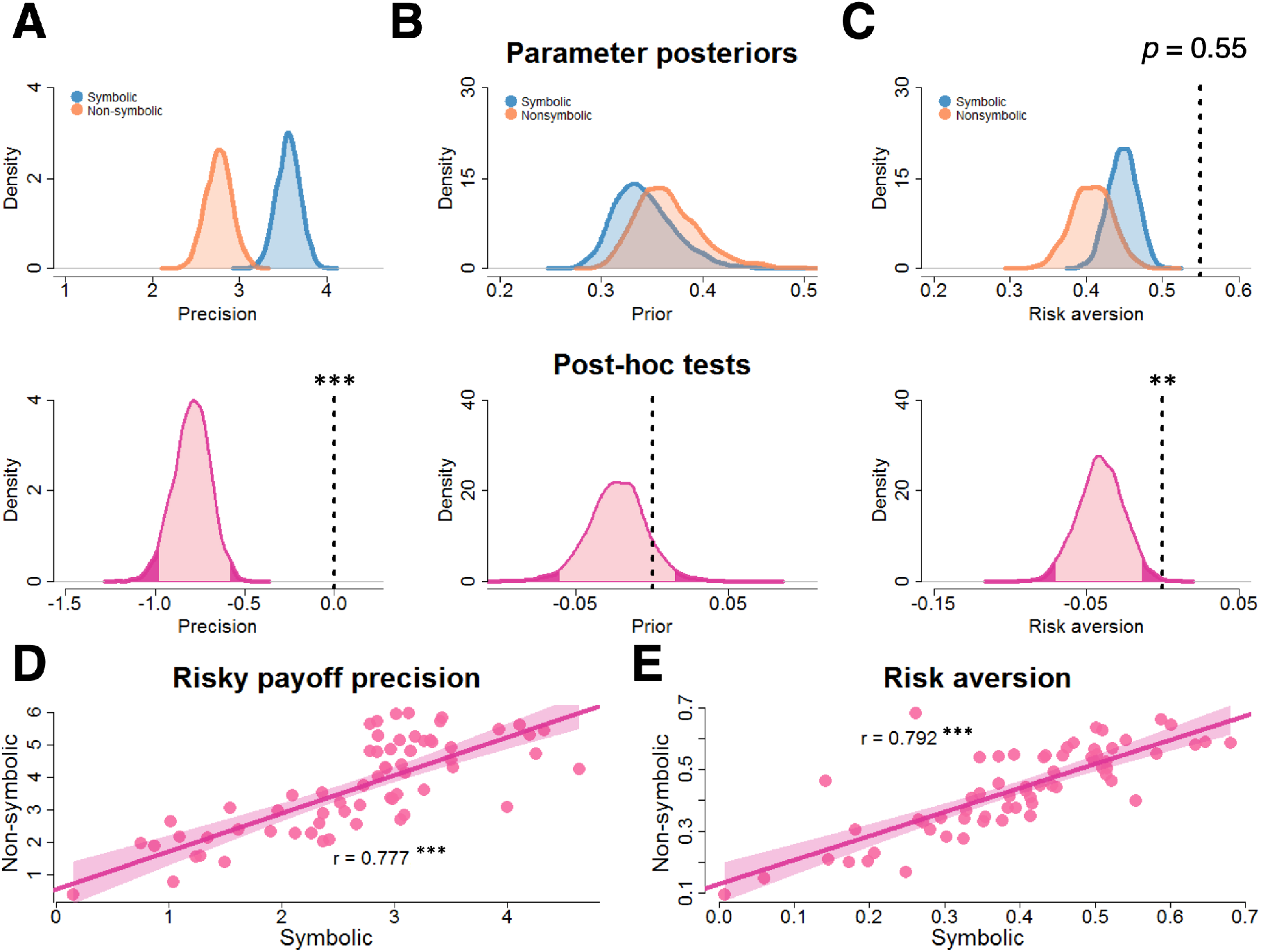
Noise in mental magnitude representations is individually robust across different types of visual displays of risky prospects. Population posterior distributions of **(a)** risky payoff precision, **(b)** the prior, and **(c)** risk aversion for risky symbolic payoffs and risky non-symbolic payoffs in risky choice behaviour. Top plots are distributions for both types of displays (symbolic payoffs in blue, non-symbolic payoffs in yellow) while bottom plots are distributions of differences between display formats (in pink). The light-shaded mass of the highest density interval (HDI) covers 95% of the posterior distribution while the dark-shaded tail-ends represent 5% of the posterior distribution. In top plots, the vertical dashed line represents the ‘rational’ risk-neutral probability, *p* = 0.55, whereas in bottom plots, it represents zero. Representations were more precise during symbolic payoffs, *γ*_*symbolic*_, than non-symbolic payoffs, *γ*_*nonsymbolic*_. This is reflected as larger risk aversion (or lower risk-neutral probability) in non-symbolic payoffs, *π*_*nonsymbolic*_ relative to symbolic payoffs, *π*_*symbolic*._ There is no difference between the priors, *σ*_*nonsymbolic*_ and *σ*_*symbolic*_. * *p* < 0.05, ** *p* < 0.01, and *** *p* < 0.001. Individual measures of **(d)** risky payoff precision and **(e)** risk aversion for symbolic and non-symbolic payoffs are related: the two measures are positively correlated. * *p* < 0.05, ** *p* < 0.01, and *** *p* < 0.001.

In line with our second hypothesis – that common magnitude representations are employed during both display types, but are subject to differing sensory noise - we found significant positive correlations across visual display types between the risk precision measures (**Fig 4d**, *r*_*γ*_ = 0.777, *p* < 0.001), the measures for the indifference point (**Supplementary Fig 2e**, *r*_*γ*_ = 0.703, *p* < 0.001), and the risk-neutral probabilities (**Fig 4e**, *r*_*π*_ = 0.792, *p* < 0.001). Finally, we plotted risk-neutral probability, *π*, as a function of precision, *γ*, for all visual displays (**Fig 5a**). The significant positive correlations between risk-neutral probability, *π*, and precision, *γ*, across all visual displays (*r*_*nonsymbolic*_ = 0.734, *p* < 0.001; *r*_*symbolic*_ = 0.567, *p* < 0.001) show that individual risk attitudes are indeed systematically related to the precision of mental magnitude representations as predicted by the model, and that this precision may reflect a psychophysical trait that is robust to sensory noise inherent in specific visual displays.

**FIGURE 5.**
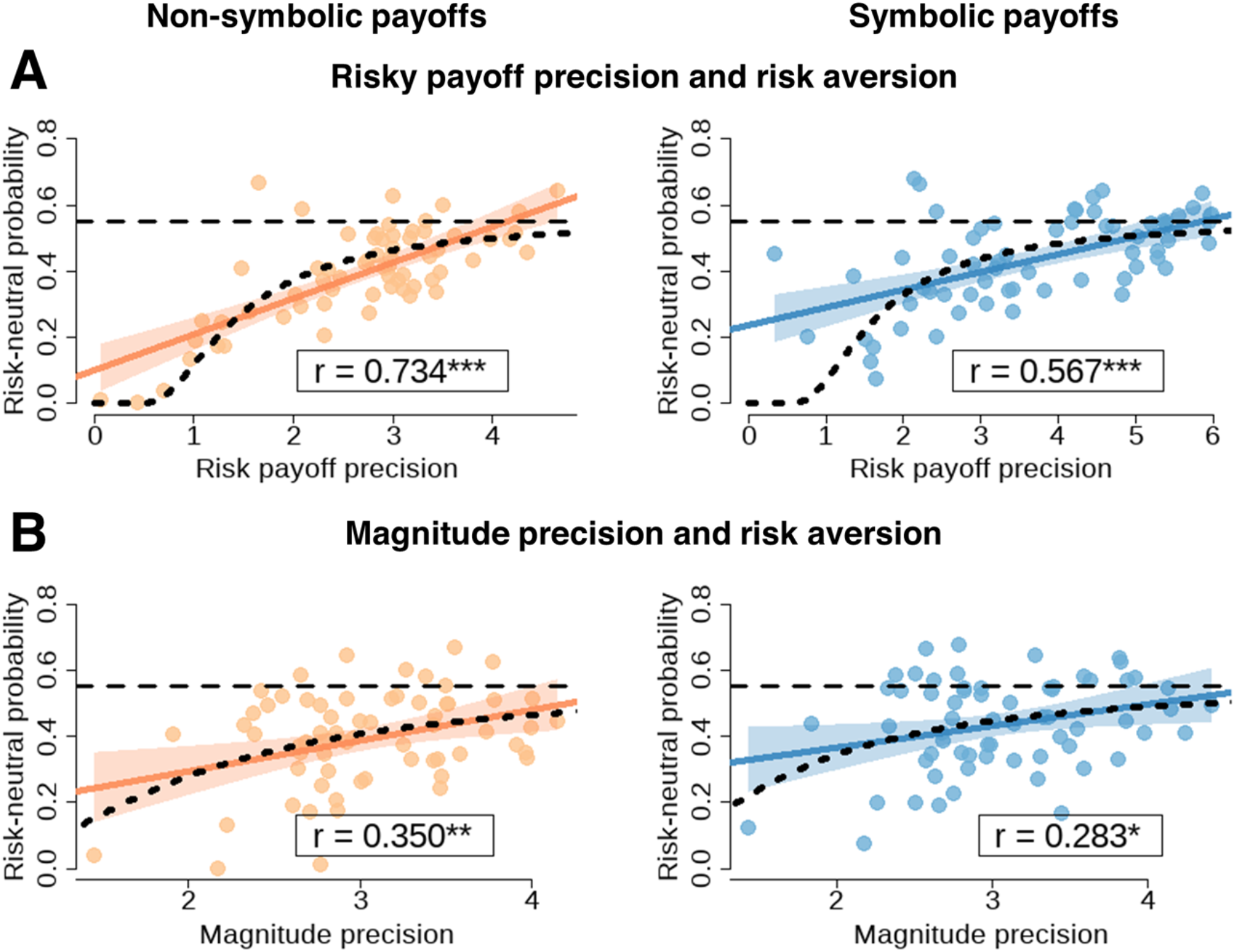
The precision of mental magnitude representations systematically relates to individual risk attitudes. **(a)** The estimated precision of the representation of potential payoffs, *γ*_*nonsymbolic*_ and *γ*_*symbolic*_, and the index of risk-aversion (measured by individual risk-neutral probability, *π*_*nonsymbolic*_ and *π*_*symbolic*_) are related across all visual displays. The curved dashed line is the prediction from the psychometric model linking the relationship between risk payoff precision and risk-neutral probability. Subjects (represented as circular dots) whose risk neutral probability are on the horizontal dashed line are risk-neutral while subjects below the dashed line are risk-averse and subjects above the dashed line are risk-seeking. **(b)** The estimated precision of magnitude representations during perceptual decision-making and individual risk attitudes are related, consistently across visual display type. The horizontal dashed lines represent risk-neutral behaviour while the curved dashed line is now the prediction linking risk-neutral probability with magnitude comparison precision. *p-*values were estimated from Pearson correlations: * *p* < 0.05, ** *p* < 0.01, and *** *p* < 0.001.

### Risk aversion in financial choices correlates with precision in mental magnitude representation during independent perceptual tasks

We tested whether the individual precision of mental magnitude representations generalizes across both perceptual and risky decision-making tasks, and whether risk aversion in the financial choices can thus be predicted by the preceding, fully independent perceptual task. We first related the slope of the psychometric curves of the independent perceptual magnitude decision-making task, *γ*_*perceptual*_ to the analogous ‘consistency’ slopes from the two visual display formats, *γ*_*nonsymbolic*_ and *γ*_*symbolic*_ of the risky-choice task, across subjects (**Supplementary Fig 3a**). In line with a shared representational mechanism, we found significant positive correlations for both display formats (*r*_*nonsymbolic*_ =0.349, *p* = 0.0025; *r*_*symbolic*_ = 0.437, *p* < 0.001). Second, we tested whether individual apparent risk aversion was systematically related to the precision of mental magnitude representations from the separate perceptual task. Indeed, we found significant positive correlations between our measure of precision from the perceptual magnitude task, *γ*_*perceptual*,_ and apparent risk aversion for the financial choices, in both presentation formats, *π*_*nonsymbolic*_ and *π*_*symbolic*_ (**Fig 5b**, *r*_*nonsymbolic*_ =0.350, *p* = 0.0025; *r*_*symbolic*_ = 0.283, *p* = 0.012). This provides crucial evidence that common mental magnitude representations are used as basis for both perceptual and economic choices, and that seemingly irrational biases in economic choice may in fact be rooted in basic properties of perceptual magnitude representations.

### Risk aversion is related to the precision of neural magnitude representations

Our findings so far suggest that both perceptual and risky financial choices are determined by the noisiness of domain-general magnitude representations that are similarly employed across different types of tasks and visual displays. We finally tested to what degree this noise and the ensuing risk aversion in financial choices is also related across purely behavioural and neural measurement techniques, as already shown in the perceptual domain (Kersey and Cantlon, 2017; Lasne et al., 2019). Such a relation would entail that just by measuring the noise in perceptual neural magnitude representations with fMRI, one would already gain information about an individual’s risk aversion in future financial choices. To test this hypothesis, we first examined whether the precision of risky choice behaviour, *γ*_*risk*_, and apparent risk aversion, *π*, were correlated with neural measures of representational precision in the perceptual task. This showed that the risk precision, *γ*_*risk*_, correlated only weakly with the *neural precision* measure for either display (*r*_*nonsymbolic*_ = 0.12, *p* = 0.17; *r*_*symbolic*_ = 0.12, *p* = 0.18; **Supplementary Fig 3b)**, but correlated significantly with *neural diminishing sensitivity* for non-symbolic payoffs (*r*_*nonsymbolic*_ = −0.21, *p* = 0.045; **Supplementary Fig 3c**, *orange markers*) and marginally for symbolic payoffs (*r*_*symbolic*_ = −0.18, *p* = 0.08; **Supplementary Fig 3c**, *blue markers*). Correspondingly, the risk-neutral probability *π* correlated significantly with *neural precision* for non-symbolic payoffs (*r* = 0.26, *p* = 0.02; **Fig 6a**, *orange markers*) and showed a non-significant relation in the same direction for symbolic payoffs (*r* = 0.09, *p* = 0.25; **Fig 6a**, *blue markers*). A similar pattern was evident for *neural diminishing sensitivity*, which correlated significantly with *π* (*r* = −0.27, *p* = 0.02; **Fig. 6b**, *orange markers*) for non-symbolic payoffs and non-significantly for symbolic payoffs (*r* = −0.14, *p* = 0.13; **Fig 6b**, *blue markers*). Thus, we found evidence, albeit less robust across visual display types, that the noisier the neural magnitude representations (*neural precision*) and the stronger the deterioration of neural representation for larger magnitudes (*neural diminishing sensitivity*), the more risk averse the individual.

**FIGURE 6.**
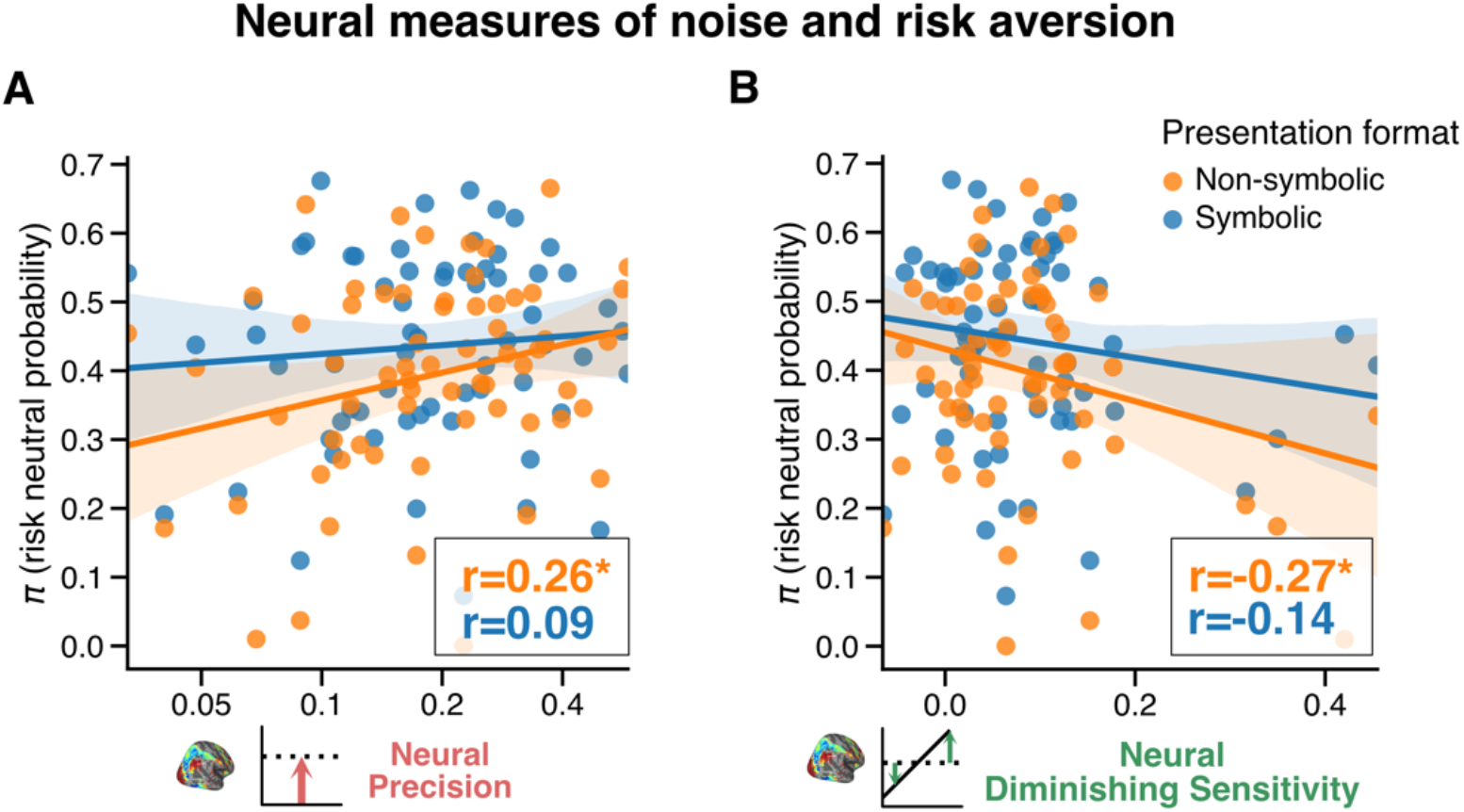
Relationship between neural measures of the precision of magnitude representations and risk aversion. **(a)** Subjects with *more precise neural magnitude representations* tended to be *less risk averse* for the separate financial choices (in particular for non-symbolic presentation format). **(b)** Similarly, subjects who showed larger *neural diminishing sensitivity* tended to be more risk averse. Both findings are in line with the behavioural finding that more risk-averse people perform better on the purely perceptual numerosity task. * *p* < 0.05.

### Effects of neural magnitude precision on individual risk aversion are mediated by noise in mental magnitude representations

While the purely behavioural estimates of magnitude precision were strongly related across all task and display types, the neural and behavioural magnitude representation measures were more strongly related within the perceptual task than across both choice types. This suggest that non-specific noise during the perceptual fMRI measurements (e.g., measurement, physiological) may have overshadowed the relationship between neural magnitude precision and risk aversion measured outside the scanner. To account for all our measures using a single integrative framework, we thus performed mediation analyses to test whether risk aversion related specifically to that part of the variance in *neural precision* that was correlated with *behavioural* precision. Indeed, behavioural precision, *γ*_*perceptual*,_ significantly mediated (*α* × *β*) the effect between *neural precision* and risk aversion, *π*, for both non-symbolic (*p*_*mcmc*_ = 0.013) and symbolic (*p*_*mcmc*_ = 0.019) visual displays (**Fig 7a,b**). In line with this result, *γ*_*perceptual*_ also significantly mediated the effect between neural precision and risky payoff precision for both symbolic (*γ*_*nonsymbolic*_; *p*_*mcmc*_ = 0.008) and symbolic (*γ*_*nonsymbolic*_; *p*_*mcmc*_ = 0.004) displays (**Supplementary Fig. 4a,b**).

**FIGURE 7.**
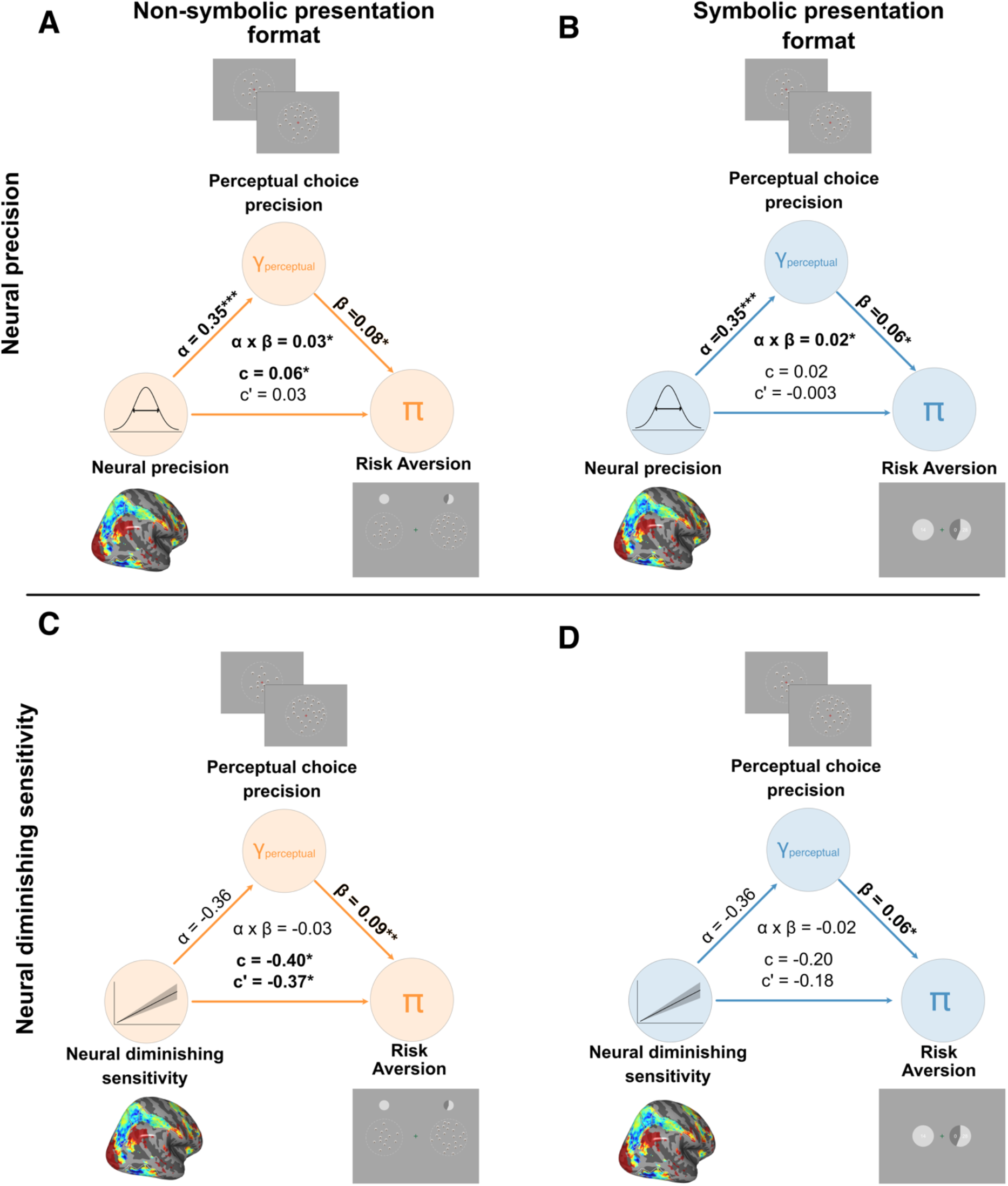
The mediating effect of behavioural magnitude precision reveals the association between neural magnitude precision and risk aversion. The effect of decoded neural precision on risk aversion is significantly *mediated* by *γ*_*perceptual*_ for **(a)** no-symbolic and **(b)** symbolic presentation formats. In contrast, there is a *direct effect* of neural diminishing sensitivity on risk aversion for **(c)** non-symbolic payoffs and no mediation effect from *γ*_*perceptual*._ This effect was not significant for **(d)** symbolic payoffs (mirroring the correlation pattern in Fig 6b). Bayesian “*p-*values” were calculated using hierarchical Bayesian mediation analysis (see **Supplementary Fig. 5a**): * *p* < 0.05, ** *p* < 0.01, and *** *p* < 0.001.

On the other hand, for *neural diminishing sensitivity* we found no such mediating effect of *γ*_*perceptual*_ on *π* and *γ*_*nonsymbolic*_, *γ*_*symbolic*_, but a significant *direct* effect (*c*′) on *π* for non-symbolic payoffs (*p*_*mcmc*_ = 0.017; but no effect for symbolic payoffs, *p*_*mcmc*_ = 0.16) (**Fig 7c,d**) and marginal effects on *γ*_*nonsymbolic*_ (*p*_*mcmc*_ = 0.054) and *γ*_*symbolic*_ (*p*_*mcmc*_ = 0.08) (see also **Supplementary Fig 4**). These findings are in line with the significant positive relationship between *neural diminishing sensitivity* and risk aversion for non-symbolic payoffs (see above), which suggests that diminishing sensitivity may capture a latent feature of neural processing that is not fully accounted for by the NLC model. Irrespective of these details, our data confirm that neural precision is an aspect of the same latent trait that determines the precision of mental magnitude representations, and that this trait is related to risk aversion after controlling for unspecific noise.

## DISCUSSION

Individual differences in risk aversion have traditionally been thought to emerge from valuation processes, either as a consequence of individual differences in the concavity of the utility curve (Holt and Laury, 2002; Rabin and Thaler, 2001) or as individual ‘appetites’ for outcome variability (Gai and Vause, 2004; Hertwig et al., 2019). Here we provide direct behavioural and neural evidence to support a perceptual account of risk aversion (Frydman and Jin, 2022; Khaw et al., 2021) proposing that, at least in certain contexts, risk aversion may also arise from systematic biases of perceiving potential payoffs. A crucial prediction of this account is that the degree of risk aversion should be related to the fidelity with which subjects perceive numerical magnitudes. In line with this prediction, we found that both behavioural performance during fast, intuitive numerosity judgements, as well as the fidelity with which the corresponding magnitudes could be read-out from neural fMRI activity, are mechanistically related to risk aversion in independent risky choices that were taken under very different circumstances. Our data thus suggest that risk-averse decision-makers may not actually “shy away from risks” - instead, they may attempt to rationally maximise expected reward, but may be limited in their ability to do so because of cognitive limitations that make them underestimate the larger payoff magnitudes that come with increased risk, just as they also underestimate larger magnitudes in purely perceptual tasks.

Previous work has already sought to understand the link between numeracy and rationality in economic choice (Kacelnik and Brito e Abreu, 1998; Schley and Peters, 2014) and financial decision-making (Frydman and Jin, 2022; Khaw et al., 2021), but purely on the basis of behavioural data, without any study of the underlying neurocomputational processes. By applying a unifying, comprehensive computational model rooted in normative theories from perceptual computational neuroscience (Wei and Stocker, 2015; Woodford, 2020), we show here how risk aversion may mechanistically emerge from the precision of neural and mental magnitude representations. These mechanistic links are domain-general not only across differences in visual displays with different sensory noise but also across tasks with different behavioural goals (perceptual accuracy versus maximising financial value). Our results thus provide evidence that a similar neurocognitive mechanism guides both numerical magnitude perception and risky choice, suggesting that at least part of the individual differences in risk-averse behaviour can be explained by variability in the acuity of numerical magnitude perception.

Our results contribute to an ongoing research program that seeks to understand the extent to which principles of lower-level perceptual neural processing can account for idiosyncrasies in economic and risky decision making (Khaw et al., 2021; Louie et al., 2013; Polanía et al., 2019; Rustichini et al., 2017; Summerfield and Tsetsos, 2012; Woodford, 2020). Previous neuroimaging work along these lines has so far mainly focussed on characterising how neural valuation processes may be constrained by such principles (Grueschow et al., 2015; Polanía et al., 2014), and investigations of perceptual magnitude coding in parietal cortex had largely been confined to perceptual tasks (Dehaene, 2003; Harvey et al., 2013; Merten and Nieder, 2009; Nieder and Dehaene, 2009). However, work in psychophysics (Brus et al., 2019; Dehaene, 2003; Pardo-Vazquez et al., 2019; Weber, 2004) and on efficient (Wei and Stocker, 2015, 2017) and noisy (Petzschner et al., 2015; Pouget et al., 2013, 2016; Stocker and Simoncelli, 2006) sensory coding has proposed that seemingly fundamental properties of perceptual processing (e.g., Weber’s law or regressive bias) may in fact generalise across many task contexts. Recent work has also shown that neurons in inferior parietal sulcus are tuned to represent magnitudes beyond pure numerosities, but more generally across multiple domains, such as time intervals (Harvey et al., 2020) and object sizes (Harvey et al., 2015). This makes it plausible that such psychophysically-defined perceptual limitations may also affect behaviour in higher-order cognitive domains, as we show here for risk-taking in the financial domain.

That being said, our results do not imply that individual variability in risky choice solely reflects properties of parietal-encoded mental magnitude representations. Previous studies have identified other sources of neural variability that may relate to individual differences in risky decision-making, such as fluctuations in striatal regions (Blankenstein et al., 2018; Chew et al., 2019) and in anterior insula (Paulus et al., 2003; Preuschoff et al., 2008) or even neural lateralisation (Sacré et al., 2019). More generally, recent findings have also suggested that economic choice variability may be associated with weaker value signals in orbitofrontal cortex (Shi et al., 2022). Thus, our results highlight the precision of parietal magnitude representations as just one of several fundamental sources of decision-relevant noise that can lead to individual choice variability and risk attitudes. In line with the hypothesis of value construction (O’Doherty et al., 2021; Shi et al., 2022), the brain may flexibly and actively construct subjective value representations from several context-dependent attributes, with noise in magnitude representations being one of the brain features affecting economic choice variability and bias, alongside more affective reward- (Chew et al., 2019) and value-based (Conen and Padoa-Schioppa, 2015; Padoa-Schioppa, 2013; Padoa-Schioppa and Conen, 2017) neural variability. The degree to which individual or contextual choice variability and bias reflects mixtures of perceptual-, reward-, value-based, or even emotional factors (Kusev et al., 2017; Paulus et al., 2003) is thus an exciting question that should be considered in future work.

Our findings that perceptual and neural magnitude noise can mechanistically affect economic choice bias and variability suggest that normative and predictive models such as the NLC can capture basic magnitude representations commonly underlying both risky choice as well as perceptual judgments. However, the current form of the NLC is actually agnostic to whether choices are information-maximising (i.e., perceptual choice) or maximising expected reward or payoff (i.e., economic choice). Previous results have suggested that models can be set up to explicitly dissociate these two behavioural goals (Heng et al., 2020; Park and Pillow, 2020; Rustichini et al., 2017; Schaffner et al., 2021). However, while the NLC may not be as computationally detailed as these more recent optimal coding models, our approach establishes it as one of the few choice models for whom there is an empirical correspondence between model parameters and independent measures of neural processing. This paves the way for other uses of this general approach to validate complex model assumptions, and to predict choice behaviours based on independent measures of basic neural processes.

Our approach of demonstrating that risk-averse behaviour is partially rooted in capacity constraints of perceptual brain processes dovetails with other (neuro)economic choice models that have taken inspiration in neurocomputational accounts of vision (Khaw et al., 2017; Louie et al., 2013), sensory processing (Heng et al., 2020; Polanía et al., 2019), perception (Fiedler and Glöckner, 2012; Frydman and Jin, 2022), attention (Gluth et al., 2018; Krajbich et al., 2010), or memory (Azeredo da Silveira and Woodford, 2019), among others.

Crucially, our specific neurocognitive account of risk aversion illustrates how a principled understanding and empirical measures of basic brain mechanisms may lend credence to choice models that have originally been developed mainly based on theory and/or fits to empirical choice data. Thus, perhaps in analogy to how economic modelling of choice data may benefit from being constrained by choice axioms (Caplin and Dean, 2009), our study suggests that the vast space of possible choice models can be narrowed down by empirical measures of the basic information-processing operations assumed by the model. Our approach therefore directly runs counter to previous concerns that the study of brain processes and neural data may provide little information of relevance to knowledge and theories about economic choice (Gul and Pesendorfer, 2008). In fact, more recent economic models have already begun to propose cognitive micro-foundations that offer interesting computational hypotheses about choice variability, bias, and context-dependent behaviour, e.g., (Bordalo et al., 2012, 2019; Frydman and Jin, 2022; Khaw et al., 2021; Steverson et al., 2019; Webb, 2019). However, we highlight again that cognitive or economic models that are theoretically based on a neurobiological account should substantiate their assumptions with neural data. This will be essential for disambiguating between the ballooning number of choice models that draw on such theoretical frameworks (Bhatia et al., 2021; He et al., 2022), and to establish which of the assumptions underlying these flexible models are in fact plausible in the light of empirical data.

Finally, our current results offer correlational evidence of a relationship between mental magnitude representations encoded in parietal cortex and risk-averse behaviour as predicted by the NLC model. More powerful future tests of the model may investigate with brain stimulation methods whether parietal cortex is indeed causally involved during risky decision-making, and to what degree a perceptual account of risky-choice behaviour can be generalised to atypical populations, such as patients diagnosed with dyscalculia (Butterworth et al., 2011; Price et al., 2007), impulse-control disorders (Specker et al., 1995), and chronic stress (Engelmann et al., 2015). It would also be interesting to map the correspondence between numerical ability and risky choice behaviour across various stages of human development with dynamic changes of risk attitudes (Tymula et al., 2013); or even across various real-world contexts with strong differences in risk attitudes, such as in countries with different stages of economic development (Dillon et al., 2017; Haushofer and Fehr, 2014). Last but not least, our findings may also have policy implications if we could reliably measure such representations in environmental contexts that go beyond controlled laboratory settings. For example, studies on educational outcomes have shown that increasing numeracy has clear long-term consequences in improving financial literacy and lifelong incomes, which may depend on the individual’s ability to accurately gauge and evaluate risk (Almenberg and Widmark, 2011; Peters, 2020; Skagerlund et al., 2018).

## MATERIALS AND METHODS

### Participants

Sixty-four right-handed participants (26 females, ages 18 to 35) volunteered to participate in this study. We informed them about the study’s objectives, the equipment used in the experiment, the data recorded and obtained from them, the tasks involved, and their expected payoffs. We also screened participants for MR compatibility prior to their participation in the study. No participant had indications of psychiatric or neurological disorders or needed visual correction. Our experiments conformed to the Declaration of Helsinki and our protocol had the approval from the Canton of Zurich’s Ethics Committee.

### Procedure

Participants completed the MRI screening and consent forms upon their arrival, before the experiment. They then went into a behavioural testing room and read the instructions for the perceptual magnitude and risky choice tasks, as well as information on MRI safety. Participants performed two tasks sequentially: a perceptual magnitude task and an economic risky-choice task. Participants first completed the perceptual magnitude task inside the MRI scanner, where we recorded their behavioural and neural measures of mental magnitude precision. Participants then subsequently completed the risky choice task outside the MR scanner, inside a behavioural testing room. We designed the experiment in a way that they first performed the perceptual magnitude task before the gambling task, to prevent the statistics of the gambling task from altering participants’ priors and thereby influence the neural measures of magnitude representation. We also recorded and collected peripheral pupil and physiological measures, particularly eye movements, heartbeat and breathing measurements during the perceptual magnitude task while we recorded and collected behavioural measures of magnitude precision. After completing both tasks, we paid participants based on both their cumulative score in the perceptual magnitude task and one decision trial randomly selected by our algorithm in the risky choice task (see below). We additionally paid out a show-up fee of 10 CHF for their attendance and participation. Participants familiarised themselves with the tasks and performed practice trials of both before they were brought to the MRI scanner room.

### Perceptual magnitude task

Participants had to choose which of two sequentially presented clouds of coins contained a larger quantity of coins. Before the start of every trial, a red fixation cross was presented for 1 second. Then the first cloud, *m*, was overlaid on the red fixation for 600 milliseconds. After an interval lasting between 6 and 9 seconds the second set of coin clouds, *n*, appeared on the screen. Only the red fixation remained on the screen during this interval. We chose the presentation timing and interval length in a way that would provide sufficient time to model the haemodynamic response function (HRF) from neural data and would prevent the HRF response of the first stimulus presentation from being contaminated by response-related activity during the second stimulus presentation. The second set appeared for another 600 milliseconds, whereafter the fixation cross changed from red to green, prompting participants to decide which cloud had the larger quantity of coins. They had 2.5 seconds to respond. A green-coloured letter “*l*” appeared on the screen to indicate participants had pressed left and if they chose the first set; otherwise, when participants pressed right and had chosen the second cloud, a green-coloured letter “*r*” appeared on the screen. Responses made too early (fixation had not turned green) or too late (after 2.5 seconds and when fixation had reverted back to red) were labelled as missed responses. Each correct response corresponded to a reward of 0.25 CHF, but participants had no feedback on the accuracy of their responses or the accumulation of points throughout the task. The perceptual magnitude task had a total of 216 trials distributed across 6 runs and lasted a total of 30 to 40 minutes. The first set varied from 5 to 28 were drawn from a geometric sequence with steps of 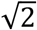(5, 7, 10, 14, 20, 18), while we varied the second set by multiplying each magnitude from the first set by a factor of 2^*h*/4^ where *h* is in discrete steps from −6 to 6.

### Risky choice task

Participants had to choose between a certain gamble with monetary offer, *s*, and a 100% probability of payout, and a risky gamble with a monetary offer, *r*. We fixed the probability of payout for the risky offer at *p* = 0.55. Thus, a participant choosing the risky gamble had a 55% chance of being paid the risky offer and a 45% chance of no payment. When a participant chose the safe option, they had a 100 percent probability of payout. During the task, the monetary payouts were presented in two display formats: symbolic payoffs of Arabic numerals or a cloud of 1-CHF coins. The gambles were presented simultaneously at the left and right sides of the monitor screen. In the beginning of every trial, the stated certain and risky probabilities appeared on the screen alongside the red fixation cross, with the position (left/right from the fixation cross) of these probabilities varying randomly from trial to trial. We used grey-shaded pies to represent the probabilities in both formats. The light-grey shade in the risky gamble represented the probability the individual would receive the stated monetary amount, while the dark grey shade represented the probability the individual would receive nothing. We overlaid numerical monetary offers inside the probability pies when we displayed them as risky symbolic payoffs while we positioned the amounts below the pies when displayed as a cloud of 1-CHF coins. Both display formats were presented in alternating blocks of 40 trials per block, totalling 12 blocks. The monetary amounts were displayed once the fixation cross changed from red to green, and participants had 3 seconds to choose the gamble on the left or on the right. A green-coloured letter “*l*” appeared on the screen when participants pressed left and a green-coloured letter “*r*” appeared on the screen when participants pressed right. Responses made too early (fixation had not turned green) or too late (fixation had already reverted back to red) were missed responses, and these missed trials were not included in the draw of the final trial that determined the monetary payout of the participant.

After participants finished all the trials, one trial was randomly drawn as basis for the payout. If on that trial a participant chose the certain option, she would immediately receive that amount. If, on the other hand, a risky option was chosen, they had to roll from a virtual, hundred-sided die, and if the resulted roll was a number smaller than or equal to 55, they were paid out according to the indicated amount; otherwise, they received nothing. The task had a total of 480 trials lasting between 30 and 40 minutes (240 trials per display presentation format). We varied the distribution of monetary payoffs with the sure gamble varying from 5 to 28 drawn from a geometric sequence with steps of 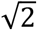similar to the perceptual magnitude task and the probabilistic lotteries varying by a factor of 2^*h*/4^ in steps of 0 to 8.

### The noisy logarithmic coding model

The NLC model assumes that the coding of numerical magnitudes occurs in logarithmic space, and that the amount of noise of this representation is constant after accounting for the log-ratio of magnitudes (i.e., *scale invariance*). The NLC prescribes that the psychometric curves of each stimulus magnitude should have similar slopes when plotted on a logarithmic scale, while a single psychometric curve can fit all the choice data when plotting the log-ratio of these magnitudes. To this end, we separately fitted six psychometric curves for the perceptual magnitude task, where each curve plots the probability of judging the second cloud of coins, *X*, to be of greater magnitude than the first cloud, *C*, as a function of the amount of *X*; and the reference of each curve is the magnitude of the first coin cloud, *C* = {5,7,10,14,20,28}. The six psychometric curves for risky choice, on the other hand, represented the probability of choosing the risky gamble, *X*, over the sure gamble, *C*, as a function of *X*; and the reference of each curve is the amount of the sure offer, *C* = {5,7,10,14,20,28}. We used a two-parameter probit model with slope, *γ*_*C*_, and intercept, *δ*_*C*_, to fit choice data in both perceptual magnitudes and risk in both linear space,

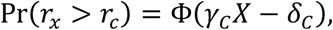

and in log space,

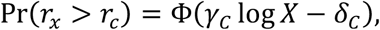

where Φ(·) is the cumulative distribution function of the standard normal distribution. We separately estimated the parameters, (*δ*_*C*_, *γ*_*C*_) at the population and individual levels using a hierarchical Bayesian framework in RJAGS (**Supplementary Fig. 5a**).

We similarly fitted the NLC using a probit model, but instead of fitting six separate psychometric curves as we did in the previous step, we fitted one psychometric curve to choice data in both tasks. The model assumes a log-ratio encoding of numerical magnitudes, and we estimated one slope, *γ*, and intercept, *δ*,

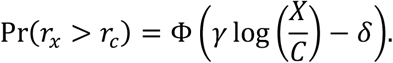

Similarly, we measured (*δ, γ*) at the population and individual levels. The NLC prescribes that *γ* measures the precision of our mental magnitude representations while *δ* contains information about the indifference point. We constrained the standard probit and rationalised it as the NLC by mapping the probit parameters with NLC model specifications,

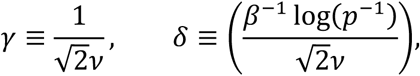

where *v* is the noise in mental magnitude representations of magnitudes; *p* is the stated probability (0 < *p* < 1 during risk and *p* = 1 during perceptual magnitudes); and, 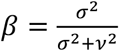is the linear weighting between the width of the prior, *σ*, and noise in mental representation, *v* (see (Khaw et al., 2021) for the full derivation).

We can then calculate an individual’s indifference point and index of risk aversion using the probit parameters. First, the indifference point is the level to which the individual is indifferent in choosing either *X* or *C*, and this indifference is determined by the following threshold rule,

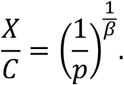

We can derive and estimate the threshold, *θ*, using the NLC’s intercept and slope parameters during risky choice (*δ*_*risk*_, *γ*_*risk*_),

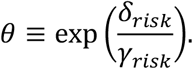

Second, we can also calculate a standard economic index of risk aversion, namely the *risk-neutral* probability: the degree to which the probability-of-payoff in the risky choice options *seems* to be underestimated. For example, if a subject is so risk-averse that her indifference point lies at 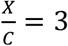 (note that an optimal, risk-neutral decision-maker would have the indifference point at 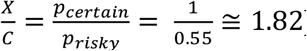), she chooses equivalent to an optimal risk-neutral decision-maker in the same paradigm but a 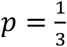. According to NLC, a decision-maker will only behave with a risk-neutral probability equal to the *objective* probability, *p* = 0.55, in the absence of noise, *v* = 0; otherwise, the individual’s risk-neutral probability in the presence of noise, *v* > 0, is,

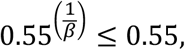

and, thus, by definition, subjects are risk averse. We can similarly derive and measure an individual’s risk-neutral probability using the NLC intercept and slope parameters (*δ*_*risk*_, *γ*_*risk*_),

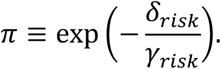

Finally, the NLC predicts a positive nonlinear relationship between the precision in mental magnitude representations, *γ*_*risk*_, and our index of risk aversion, *π*. We fitted a psychometric model that assumes a common prior, *σ* (see (Khaw et al., 2021) for more details),

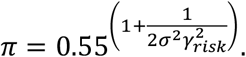

We estimated *σ* by regressing participant precision measures of payoff offers, *γ*_*risk*_, to their corresponding risk-neutral probability, *π*. We also fitted the relationship between *γ*_*risk*_ and *π* using simple linear regression. Given that our central hypothesis is to determine whether individual risk-averse behaviour can be predicted by an external measure of magnitude precision, *γ*_*perceptual*_, from an independent perceptual magnitude task, we fitted both linear and nonlinear models, but instead, regressed our index of risk aversion, *π*, on *γ*_*perceptual*_.

### Preference-based models

We also fitted behavioural data from the risky choice task using stochastic versions of standard economic models to test whether the NLC is a model that explains our empirical data better than do models that assume expected utility maximisation to explain risk aversion. We also used a hierarchical Bayesian framework to fit these models to data. We considered three classes of canonical preference-based models for model comparison, namely **(a)** constant relative risk aversion (CRRA), a standard model of expected utility theory (EUT) (Apesteguia and Ballester, 2018; von Neumann and Morgenstern, 1944); **(b)** cumulative prospect theory (CPT) (Kahneman and Tversky, 1979; Nilsson et al., 2011; Tversky and Kahneman, 1992); and, **(c)** salience theory (Bordalo et al., 2012). We used probit (i.e., the random error term, *ε*, is drawn from a Gaussian distribution) and logit (i.e., *ε* is drawn from a Gumbel or extreme-value distribution) models to account for stochasticity during risky choice.

#### Constant relative risk aversion

In CRRA, the subjective utility of the monetary offer, *y*, is represented by a utility function, *u*(·), which takes the form,

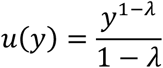

where the parameter, *λ*, accounts for the nonlinearity of subjective utility over *y*. We can then calculate the expected utility of each of the monetary amounts, *y* ∈ {*X, C*}, and then estimate the probability of choosing the risky relative to the sure gamble,

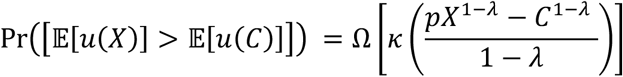

where *κ* is the sensitivity parameter and Ω(·) represents the distribution function of either Probit or Logit error distributions.

#### Cumulative Prospect theory

Distortions in monetary offers in CPT are accounted for by the value function, *v*(·),

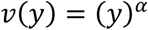

such that *α* > 0 is the parameter which accounts for the degree of risk preference (*α* < 1 suggests risk aversion; otherwise, risk-seeking behaviour); and, for completeness, *ω*(·) is the probability weighting function of the form,

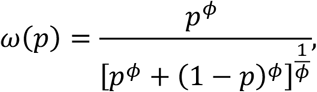

where the parameter, *ϕ*, accounts for the nonlinear probability distortion. By combining payoff and probability distortions into a weighted value function, *V*(*y*_*k*_; *p*_*k*_) = Σ_*k*_ *ω*(*p*_*k*_)*v*(*y*_*k*_) (where *k* indexes a gamble’s possible payoff outcomes), we estimate probability of choosing the risky relative to the sure gamble,

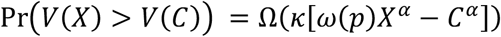

where *κ* is similarly, the sensitivity parameter, and Ω(·) is either a Probit or Logit error distribution function.

#### Salience theory

ST assumes that risky choice bias arises from salience-driven probability distortions, where these distortions are, in turn, due to both the degree of saliency and diminishing sensitivity of monetary payoffs. Both these distortionary features are captured by a salience function, *w*(·),

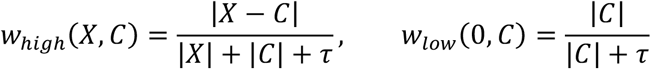

where the parameter, *τ*, accounts for the degree of diminishing sensitivity. Here, the larger the difference between the payoffs, *X* and *C*, the more salient the difference; and, the larger the *τ*, the less sensitive the decision-maker can discriminate between payoffs. Salience distorts probabilities by way of the probability distortion function, *W*(·),

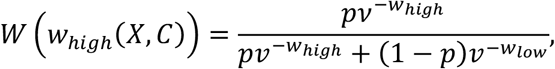

where *p* is the probability and the parameter, *v*, captures the amount of distortion from the salience function. Similarly, all of these are combined into a weighted value function, *S*(*y*_*k*_) = Σ_*k*_ *W*(*w*_*k*_)*y*_*k*_, and the probability of choosing the risky relative to the sure gamble is,

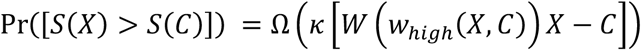

with the sensitivity parameter, *κ*, and Ω(·) is either a Probit or Logit distribution function.

### Hierarchical Bayesian parameter estimation

We estimated model parameters using hierarchical Bayesian estimation and Markov Chain Monte Carlo (MCMC) techniques (Gelman et al., 2013; Kruschke, 2015) We used a Gibbs sampler implemented in JAGS (Plummer, 2003). We used weakly informative hyperpriors for the group-level distributions. The exact model specification and used priors can be found in **Supplementary Fig. 5**. We drew a total of 50,000 burn-in samples to let the MCMC sampler reach a stationary distribution. Then, for each model, we drew a total of 50,000 new samples with three chains each. We sampled each chain using different random number generator engines and different seeds. We thinned the final sample by a factor of 50, thus resulting in a final set of 1,000 samples for each parameter. We used Gelman-Rubin tests to confirm chain convergence of each parameter. All estimated parameters in our Bayesian models showed a 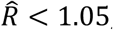, indicating that all three MCMC chains converged properly. Wherever we wanted to test whether a parameter is larger/smaller than 0, we reported Bayesian “*p*-values” that directly quantify the probability of the reported effect being smaller/larger than zero. We computed these values using posterior population distributions estimated for each parameter. During model comparison, we used the deviation information criterion (DIC) to perform model comparisons (Meyer, 2016; Pooley and Marion, 2018).

### MRI acquisition and pre-processing

We acquired functional MRI data using a Philips Achieva 3T whole-body MR scanner equipped with a 32-channel MR head coil. Specifically, we collected 6 runs with a T2*-weighted gradient-recalled echo-planar imaging (GR-EPI) sequence (189 volumes + 5 dummies; flip angle 90 degrees; *TR* = 2827 ms, *TE* = 30ms; matrix size 96 × 96, FOV 240 × 240mm; in-plane resolution of 2.5 mm; 44 slices with thickness of 2.5 mm and a slice gap of 0.5mm; SENSE acceleration in phase-encoding direction with factor 1.5; time-of-acquisition 9:14 minutes). Additionally, we acquired high-resolution T1-weighted 3D MPRAGE image (FOV: 256 × 256 × 170 mm; resolution 1 mm isotropic; *TI* = 2800 ms; 256 shots, flip angle 8 degrees; *TR* = 8.3 ms; *TE* = 3.9 ms; SENSE acceleration in left-right direction 2; time-of-acquisition 5:35 minutes) for image registration during post-processing.

Pre-processing was performed using fMRIPrep 1.4.0 (Esteban et al., 2019), which was based on Nipype 1.2.0 (Gorgolewski et al., 2011). The T1-weighted (T1w) image was corrected for intensity non-uniformity (INU) with N4BiasFieldCorrection (Tustison et al., 2010) distributed with ANTs 2.2.0 (Avants et al., 2008), and used as T1w-reference throughout the workflow. The T1w-reference was then skull-stripped with a *Nipype* implementation of the antsBrainExtraction.sh workflow (from ANTs), using OASIS30ANTs as target template. Brain tissue segmentation of cerebrospinal fluid (CSF), white-matter (WM) and grey-matter (GM) was performed on the brain-extracted T1w using fast (FSL 5.0.9) (Zhang et al., 2001). Brain surfaces were reconstructed using recon-all (FreeSurfer 6.0.1) (Fischl et al., 1999), and the brain mask that was estimated previously was refined with a custom variation of the method to reconcile ANTs-derived and FreeSurfer-derived segmentations of the cortical grey-matter of Mindboggle (Klein et al., 2017). Volume-based spatial normalization to one standard space (MNI152NLin2009cAsym) was performed through nonlinear registration with antsRegistration (ANTs 2.2.0), using brain-extracted versions of both T1w reference and the T1w template. We used *ICBM 152 Nonlinear Asymmetrical template version 2009c* (Fonov et al., 2009) as our template for spatial normalisation.

For each of the 6 BOLD runs per subject (across all tasks and sessions), we performed the following pre-processing procedure: First, a reference volume and its skull-stripped version were generated using a custom methodology of *fMRIPrep*. A deformation field to correct for susceptibility distortions was estimated based on a field map that was co-registered to the BOLD reference, using a custom workflow of *fMRIPrep* derived from D. Greve’s epidewarp.fsl <u>script</u> and further improvements of HCP Pipelines (Glasser et al., 2013). The deformation field is that resulting from co-registering the BOLD reference to the same-subject T1w-reference with its intensity inverted (Huntenburg, 2014; Wang et al., 2017).

Registration was performed with antsRegistration (ANTs 2.2.0), and the process is regularised by constraining deformation to be nonzero only along the phase-encoding direction and modulated with an average fieldmap template (Treiber et al., 2016). Based on the estimated susceptibility distortion, an unwarped BOLD reference was calculated for a more accurate co-registration with the anatomical reference. The BOLD reference was then co-registered to the T1w reference using bbregister (FreeSurfer) which implements boundary-based registration (Greve and Fischl, 2009; Jenkinson et al., 2002). Co-registration was configured with nine degrees of freedom to account for distortions remaining in the BOLD reference. Head-motion parameters with respect to the BOLD reference (transformation matrices, and six corresponding rotation and translation parameters) are estimated before any spatiotemporal filtering using mcflirt (FSL 5.0.9) (Jenkinson et al., 2002). BOLD runs were slice-time corrected using 3dTshift from AFNI 20160207 (Cox and Hyde, 1997). The BOLD time-series were resampled to surfaces on the following spaces: *fsaverage5, fsaverage6*. The BOLD time-series (including slice-timing correction when applied) were resampled onto their original, native space by applying a single, composite transform to correct for head-motion and susceptibility distortions.

These resampled BOLD time-series will be referred to as *preprocessed BOLD in original space*, or just *preprocessed BOLD*. The BOLD time-series were resampled into standard space, generating a *preprocessed BOLD run in [‘MNI152NLin2009cAsym’] space*. First, a reference volume and its skull-stripped version were generated using a custom methodology of *fMRIPrep*. Several confounding time-series were calculated based on the *preprocessed BOLD*: framewise displacement (FD), DVARS and three region-wise global signals. FD and DVARS were calculated for each functional run, both using their implementations in *Nipype* (following the definitions by (Power et al., 2014)). The three global signals were extracted within the CSF, the WM, and the whole-brain masks. Additionally, a set of physiological regressors were extracted to allow for component-based noise correction (*CompCor*) (Behzadi et al., 2007).

Principal components were estimated after high-pass filtering the *preprocessed BOLD* time-series (using a discrete cosine filter with 128s cut-off) for the two *CompCor* variants: temporal (*tCompCor*) and anatomical (*aCompCor*). *tCompCor* components were then calculated from the top 5% variable voxels within a mask covering the subcortical regions. This subcortical mask was obtained by heavily eroding the brain mask, which ensures it does not include cortical GM regions. *aCompCor* components were calculated within the intersection of the aforementioned mask and the union of CSF and WM masks calculated in T1w space, after their projection to the native space of each functional run (using the inverse BOLD-to-T1w transformation). Components are also calculated separately within the WM and CSF masks. For each CompCor decomposition, the *k* components with the largest singular values were retained, such that the retained components’ time series were sufficient to explain 50 percent of variance across the nuisance mask (CSF, WM, combined, or temporal). The remaining components were dropped from consideration. The head-motion estimates calculated in the correction step were also placed within the corresponding confounds file. The confound time series derived from head motion estimates and global signals were expanded with the inclusion of temporal derivatives and quadratic terms for each (Satterthwaite et al., 2013). Frames that exceeded a threshold of 0.5 mm FD or 1.5 standardised DVARS were annotated as motion outliers. All resamplings can be performed with *a single interpolation step* by composing all the pertinent transformations (i.e., head-motion transform matrices, susceptibility distortion correction when available, and co-registrations to anatomical and output spaces). Gridded (volumetric) resamplings were performed using antsApplyTransforms (ANTs), configured with Lanczos interpolation to minimize the smoothing effects of other kernels (Lanczos, 1964). Non-gridded (surface) resamplings were performed using *mri_vol2surf* (FreeSurfer).

Many internal operations of *fMRIPrep* use *Nilearn* 0.5.2 ((Abraham et al., 2014), RRID:SCR_001362), mostly within the functional processing workflow. For more details of the pipeline, see <u>the section corresponding to workflows in</u> *<u>fMRIPrep</u>*<u>’s documentation</u>.

### Activation modelling

After preprocessing, fMRI data of all subjects were resampled to the fsaverage6 standard surface space of Freesurfer (Fischl et al., 1999). The data were then moderately smoothed on the surface with a Gaussian kernel with a full-width-half-maximum (FWHM) of 5mm. A general linear model (GLM) was fitted to the data with one regressor per trial (36 trials per run), as well as separate regressors for all the 23 possible second stimulus arrays presented, all convolved with a canonical hemodynamic response function. This model also included: **(a)** 5 anatomical compCor components accounting for physiological noise (Behzadi et al., 2007); **(b)** 3 translational realignment parameters and their derivatives **(c)** 3 rotational realignment parameters and their derivatives (Friston et al., 1996); **(d)** DVAR and Framewise displacement time series (Power et al., 2014); **(e)** 7 discrete cosine regressors to perform high-pass filtering with a cutoff frequency of 1/128 seconds.

### Numerosity encoding model

We used a numerical population receptive field model (nPRF) *f*(*s*) (Dumoulin and Wandell, 2008), to model BOLD responses to the first stimulus array. We modelled the data separately for every vertex and for every individual, yielding thirty-six (six trial-wise regressors per stimulus type per run) activation values for each of the six possible magnitudes of the first stimulus array. We used gradient descent optimization to find a Gaussian receptive field on the logarithmic number line that best predicted number-wise beta estimates in terms of *R*^2^. The model contained four free parameters, *θ*: (a) a baseline activation, *b*; (b) a peak activation, *A*; (c) the numerical center of the logarithmic Gaussian, *μ*; and; (d) the standard deviation of the logarithmic Gaussian, *σ*. All these parameters we jointly estimated using maximum likelihood estimation.

We averaged the vertex-wise, *μ*, parameter estimates over subjects by weighting their *R*^2^ and rendered them on the *fsaverage6* cortical surface reconstruction using Pycortex. The parameter estimates were thresholded on the mean *R*^2^ across subjects at *R*^2^ > 0.09. This allowed us to qualitatively replicate the topological number fields in the parietal and frontal cortex reported by (Harvey and Dumoulin, 2017; Harvey et al., 2013) at 3-Tesla, in the group space of a large number of subjects (*n* = 64). We manually selected all vertices in and around the intraparietal sulcus (IPS) (Eger et al., 2009; Jacob and Nieder, 2009; Lasne et al., 2019; Piazza et al., 2004) that showed clear numerical sensitivity in this map. For all subjects, we used this same cortical mask, as defined is *fsaverage6*-space.

### Numerosity decoding model

We implemented a Bayesian inversion of the nPRF encoding model, *f*(*s*), extending upon previous work of encoding-decoding models (van Bergen and Jehee, 2018; van Bergen et al., 2015)^1^. This allowed us to probe the uncertainty of numerical magnitude representations, operationalized as dispersions of the posterior distributions *Pr*(*s*|*Y*), representing the probability of different numerical magnitudes, given the BOLD data of a particular trial type for a particular trial/run. We then extended the univariate encoding model, which mapped stimuli to the univariate activation patterns, *y*_*n*_ (*f*_*n*_(*s*): *s* → *y*_*n*_), with a multivariate Gaussian noise model, *ε*, yielding a conditional probability distribution over a multivariate activation pattern, *Y* = [*y*_1_, …, *y*_*n*_], given stimulus numerosity, *s*:

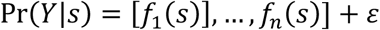

where

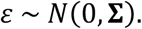

Here the covariance matrix of the noise model, **Σ**, is crucial for Bayesian decoding (van Bergen and Jehee, 2018). In our model, **Σ** is constructed as the weighted sum of (a) a perfectly-correlated covariance matrix, *ττ*^*T*^; (b) a perfectly-uncorrelated covariance matrix, *I* ∘ *ττ*^*T*^; (c) a matrix *WW*^*T*^ that quantifies the amount of overlap in receptive fields of different vertices (putatively corresponding to the overlap in neural populations); and, (d) an exponential transformation of the geodesic distance matrix on the cortical surface, exp(−*βD*):

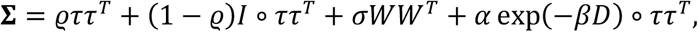

where *ϱ* ∈ [0,1] is a scalar that quantifies the noise correlation between different vertices; *τ* is a vector containing the standard deviation of the residuals of every individual vertex; *I* is the identity matrix; and, *σ*^2^ is a scalar that quantifies, in absolute terms, how much variance can be explained by overlapping neural populations.

To obtain the *W*-matrix for our nPRF model (*n* neural populations × *n* vertices) we discretized the estimated logarithmic Gaussian receptive fields into a set of two hundred logarithmically spaced bins between 1 and five times 28 (140). We can then interpret the *WW*^*T*^ matrix as the amount of overlap between the receptive fields of every vertex pair. The geodesic distance matrix, *D* represents the distance between two cortical locations on the surface, and we used it to account for spatial correlation in BOLD signal. We calculated this distance using the Heat-based method (Crane et al., 2013), as implemented in Pycortex. We modeled the proportion of this distance-based effect using the *α*-parameter, and we modeled the spatial correlation’s rate of falloff by the *β*-parameter.

We implemented the noise model in Tensorflow, used gradient descent (Kingma and Ba, 2014) to estimate *ϱ, τ, σ*^2^, *α* and *β* with a maximum likelihood cost function. We fixed 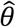 of the encoding model to the values estimated before. Fitting the noise model allows us to calculate the posterior probability of the different numerosity stimuli, *s*, conditional on the activation patterns of unseen data, *Y*^*^,

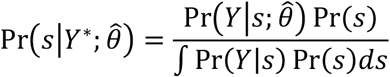

We numerically approximated the integral in 250 equally spaced steps between 1 and 50 in natural space. We used the mean of this posterior, *E*[*s*], to predict the numerosity, *s*, of the unseen data, *Y*^*^, and we used the standard deviation of the posterior to quantify the associated uncertainty of numerical representations on a trial-to-trial basis. Crucially, we fitted and evaluated the model on all data according to a leave-one-run-out cross-validation scheme, where we used the data of five runs, *Y*, for estimating the model, *f*(*s*) and *P*(*Y*|*s*), which we then used to estimate the uncertainty of numerical encoding on the sixth, left-out run, *Y*^*^.

### Model validation

To estimate the robustness of the decoding approach, we evaluated the posterior of unseen data at *p*(*s* = {5,7,10,14,20,28}) to check the mostly likely possible stimulus (maximum a posteriori; MAP stimulus) according to the model. We could then compare the accuracy of the decoding model versus a null model that would perform at chance, 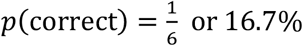.

### Individual behavioural and neural variability tests

The hierarchical Bayesian estimation procedure for both behavioural performance during perceptual magnitudes and risky choice produced posterior distributions of NLC model parameters at both population and individual levels. Thus, we extracted the mean of each subject’s posterior distribution. On the other hand, our neural encoding/decoding approach was able to extract, on a subject-to-subject basis, a measure of *neural precision*, which is the inverse of the mean standard deviation (*s*.*d*.) of the decoded trial magnitude. We also regressed these standard deviations on the stimulus magnitudes that were presented to produce a measure of *neural diminishing sensitivity*, which accounts to which extent the acuity of neural representations decreased for larger magnitudes. This measure may be less prone to general noise in the MR data that equally affects all magnitudes. The neural precision had a very non-Gaussian distribution, because some subjects had a precision very close to 0. Therefore, we log-transformed this measure before we ran any correlations. To test for individual differences, we performed simple Pearson correlation and reported correlation coefficients and corresponding p-values. We employed one-sided *p*-values to test the hypothesised relationship between individual behavioural and neural measures because we already had a very strong a priori hypothesis about the direction of possible correlations.

### Bayesian mediation analysis

We used hierarchical Bayesian mediation analysis (**Supplementary Fig. 5b**) to test whether the association between our individual neural measurements, *γ*_*neuro*_ (*neural precision* and *neural diminishing sensitivity*, obtained from our generative encoding/decoding model) in the perceptual magnitude task, and individual measurements of risk aversion, *π*_*risk*_ (and also, risk precision, *γ*_*risk*_) is mediated by individual behavioural magnitude precision, *γ*_*perceptual*_, (estimated using the NLC model). In mediation analysis (Cohen et al., 2003), we first expressed the relationship between *γ*_*risk*_ and *γ*_*neuro*_ with the following linear regression,

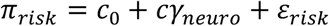

where *c*_0_ is the intercept, *ε*_*risk*_ is the residual, and the parameter *c* (the total effect) is the slope determining the strength of the relationship between *γ*_*neuro*_ and *γ*_*risk*_.

To determine whether the relationship between *γ*_*neuro*_ and *π*_*risk*_ is mediated by *γ*_*perceptual*_, we then applied structural equation modelling (SEM), which involves two sets of regression models: first, we regressed the mediating variable, *γ*_*perceptual*_ on the independent variable, *γ*_*neuro*_,

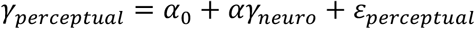

and, then we regressed *π*_*risk*_, on both independent and mediating variables

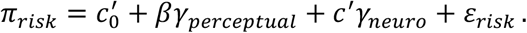

Here, 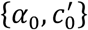 are intercepts; {*ε*_*perceptual*_,*ε*_*risk*_} are residuals; *α* measures the relationship between *γ*_*neuro*_ and *γ*_*perceptual*_; *β* measures the relationship between *γ*_*perceptual*_ and *π*_*risk*_; and, *c*′ (the *direct effect*) measures the relationship between *γ*_*neuro*_ and *π*_*risk*_ after taking into account the mediating effect of *γ*_*perceptual*_. The mediation effect, *α* × *β*, measures the relationship between *π*_*risk*_ and *γ*_*neuro*_ after accounting for the mediating effect of *γ*_*perceptual*_.

We drew inferences using a hierarchical Bayesian framework and estimated parameters using MCMC sampling in JAGS (Plummer, 2003). We used a normal distribution as our prior for the parameters for our regression coefficients and intercepts, and a Gamma distribution as our prior for the variance parameters (Miočević et al., 2018; Nuijten et al., 2015). We then estimated our Bayesian SEM using the following posterior distributions,

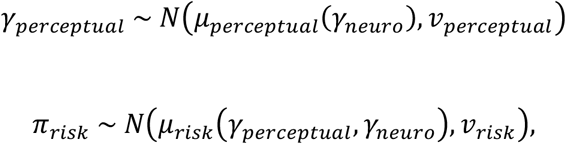

where (*v*_*perceptual*_, *v*_*risk*_) are the variances and (*μ*_*risk*_, *μ*_*perceptual*_) are the conditional means for each of the dependent and mediator variables, respectively,

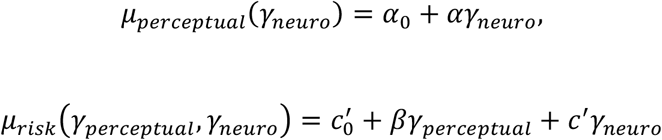

We performed posterior inferences for the mediating (*α* × *β*), direct (*c*^′^), and total (*c*) effects, for each area of numerical parietal cortex, through a Gibbs sampler implemented in RJAGS. Similar to previous analyses, we used three chains and the same initial burn-in and thinning steps to obtain a final set of 1,000 samples for each parameter at the population and individual levels. We used Gelman-Rubin tests to check whether all our latent variables had 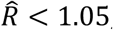, which indicated that all three MCMC chains had converged. Finally, we performed inference using Bayesian *p*-values, inferred from the highest density interval (HDI).

## SUPPLEMENTARY MATERIALS

**SUPPLEMENTARY FIGURE 1.**
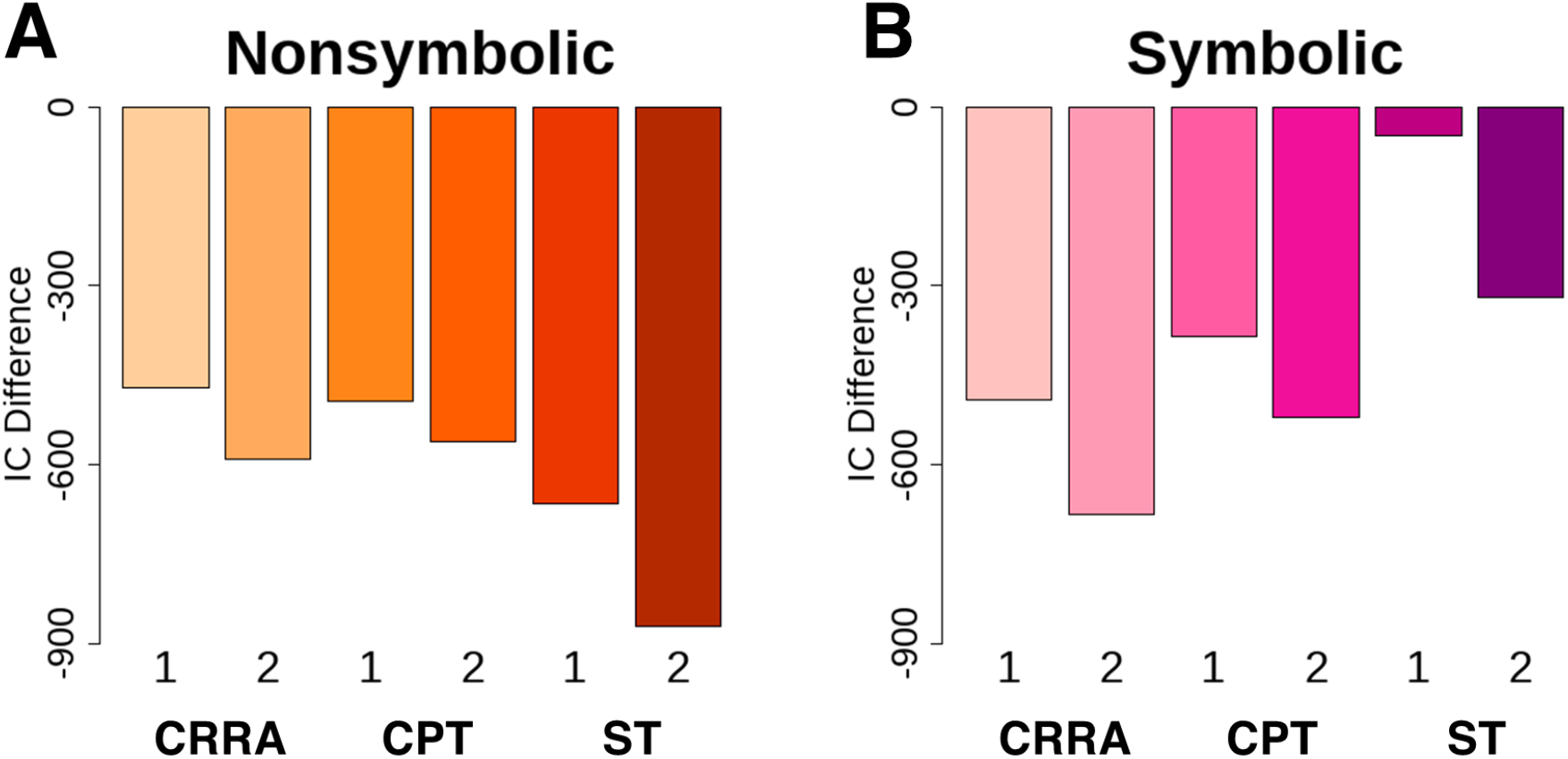
Model comparison between the NLC and competing economic choice models. DIC difference between the best model (the NLC in all cases) and the competing economic choice models, particularly constant relative risk aversion (CRRA), cumulative prospect theory (CPT), and salience theory (ST) in both **(a)** non-symbolic and **(b)** symbolic visual display formats. We fitted each of these economic choice models using the Logit (1) and Probit (2) model specifications.

**SUPPLEMENTARY FIGURE 2.**
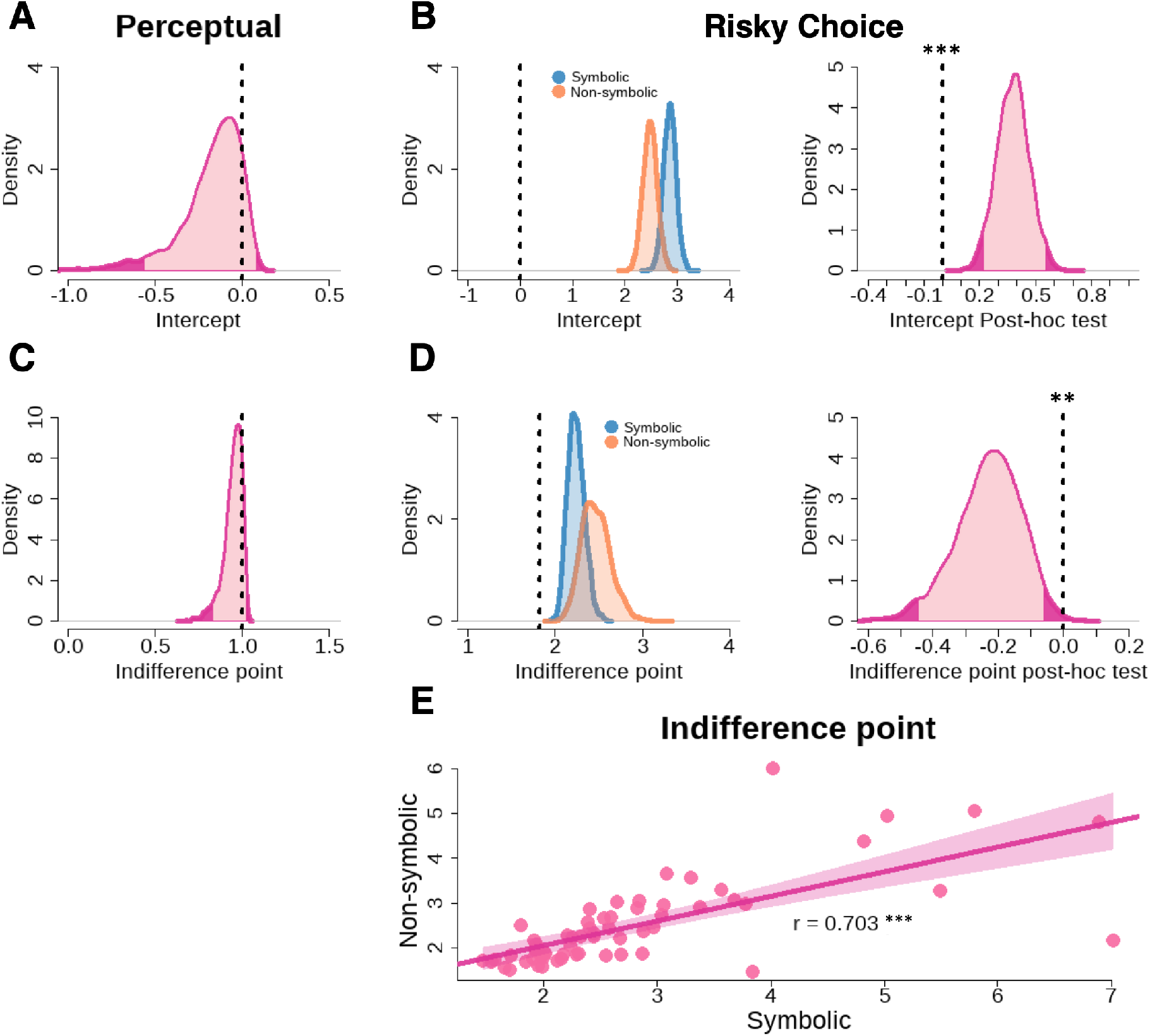
Behavioural effects of the presentation format of monetary magnitudes. **(a)** Population posterior distributions of the **(a-b)** intercept, *δ*, as well as **(c-d)** the indifference point, *θ*, for both **(a,c)** perceptual magnitude and **(b,d)** risky choice tasks. The intercept during perceptual magnitude is no different from zero (indicated here by the vertical dashed line) while it is significantly larger than zero during risky choice in both visual displays. Similarly, the indifference point in perceptual magnitude is no different from one while in risky choice, it is significantly larger than the threshold, 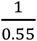 (the vertical dashed line). * *p* < 0.05, ** *p* < 0.01, and *** *p* < 0.001. **(e)** Individual measures of the indifference point for non-symbolic and symbolic payoffs are related. The two measures are positively correlated. Distributions coloured pink represent data from perceptual magnitude while blue represent data from risky symbolic payoffs and yellow-orange for risky non-symbolic payoffs. The light pink-shaded mass of the highest density interval (HDI) covers 95% of the posterior distribution while the dark-shaded tail-ends represent 5% of the posterior distribution. Post-hoc tests reveal that the posterior distribution is significantly different from zero (represented here as a vertical dashed line) if the light-shaded mass does not cross zero. * *p* < 0.05, ** *p* < 0.01, and *** *p* < 0.001.

**SUPPLEMENTARY FIGURE 3.**
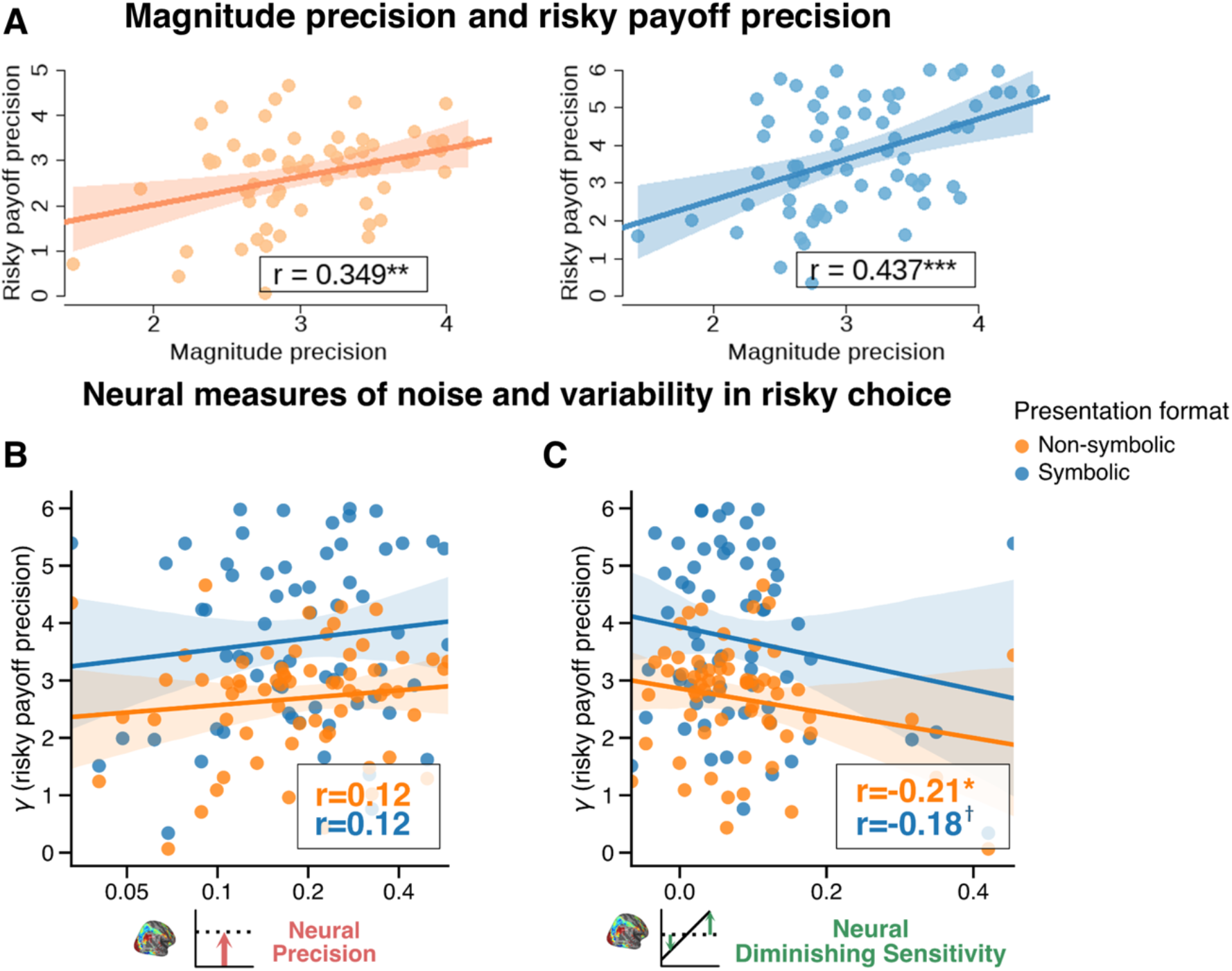
Behavioural and neural measures of representation acuity and risky choice variability. **(a)** The estimated precision of mental magnitude representations employed for the perceptual task, *γ*_*perceptual*_, and the risky decision-making task, *γ*_*nonsymbolic*_ and *γ*_*symbolic*_, are related, for both types of visual displays, as predicted by the NLC model. **(b)** The *neural precision* parameter was not correlated with the risky choice precision parameter *γ*, although the correlations were in the hypothesized direction: the higher the neural precision, the less variable the behaviour. **(c)** *Neural diminishing sensitivity* was significantly correlated with the risky choice precision parameter *γ* for the non-symbolic presentation format and marginally significant for the symbolic presentation format. *p-*values were estimated from Pearson correlations: * *p* < 0.05, ** *p* < 0.01, and *** *p* < 0.001.

**SUPPLEMENTARY FIGURE 4.**
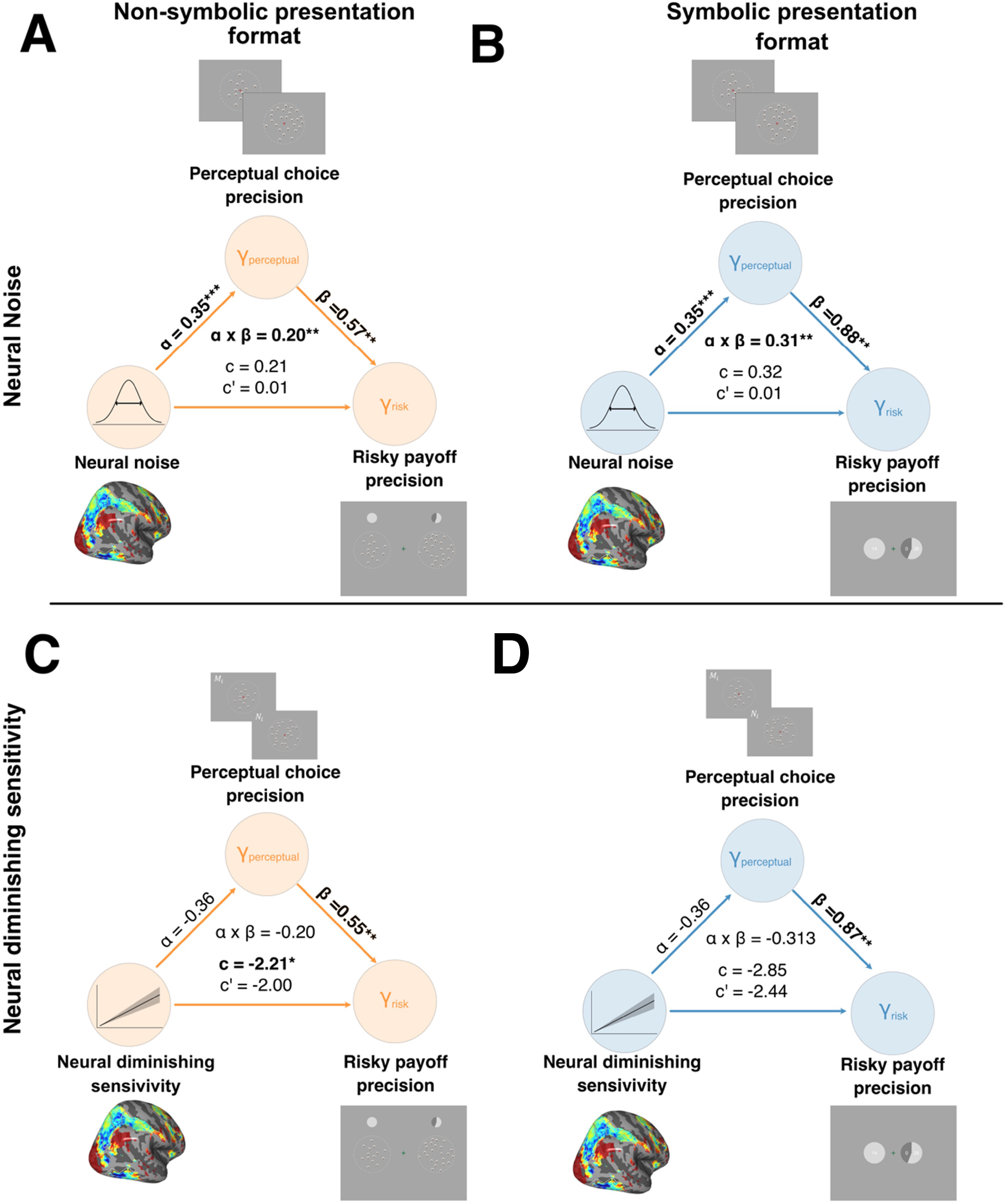
The mediating effect of behavioural precision on the association between precision-related measures on risky choice precision. **(a)** The effect of neural noise on risky choice variability for the task using symbolic numbers is mediated by perceptual choice variability. There is no significant direct or total effect. **(b)** The effect of neural noise on risky choice variability for the task using non-symbolic presentation format is mediated by perceptual choice variability. There is no significant direct or total effect. The effect of diminishing sensitivity on risky choice variability for the task using **(c)** non-symbolic numbers is mediated by perceptual choice variability, and less so with **(d)** symbolic numbers. Bayesian “*p-*values” were calculated using hierarchical Bayesian mediation analysis (see **Methods** and **Supplementary Fig 5b**): * *p* < 0.05, ** *p* < 0.01, and *** *p* < 0.001.

**SUPPLEMENTARY FIGURE 5.**
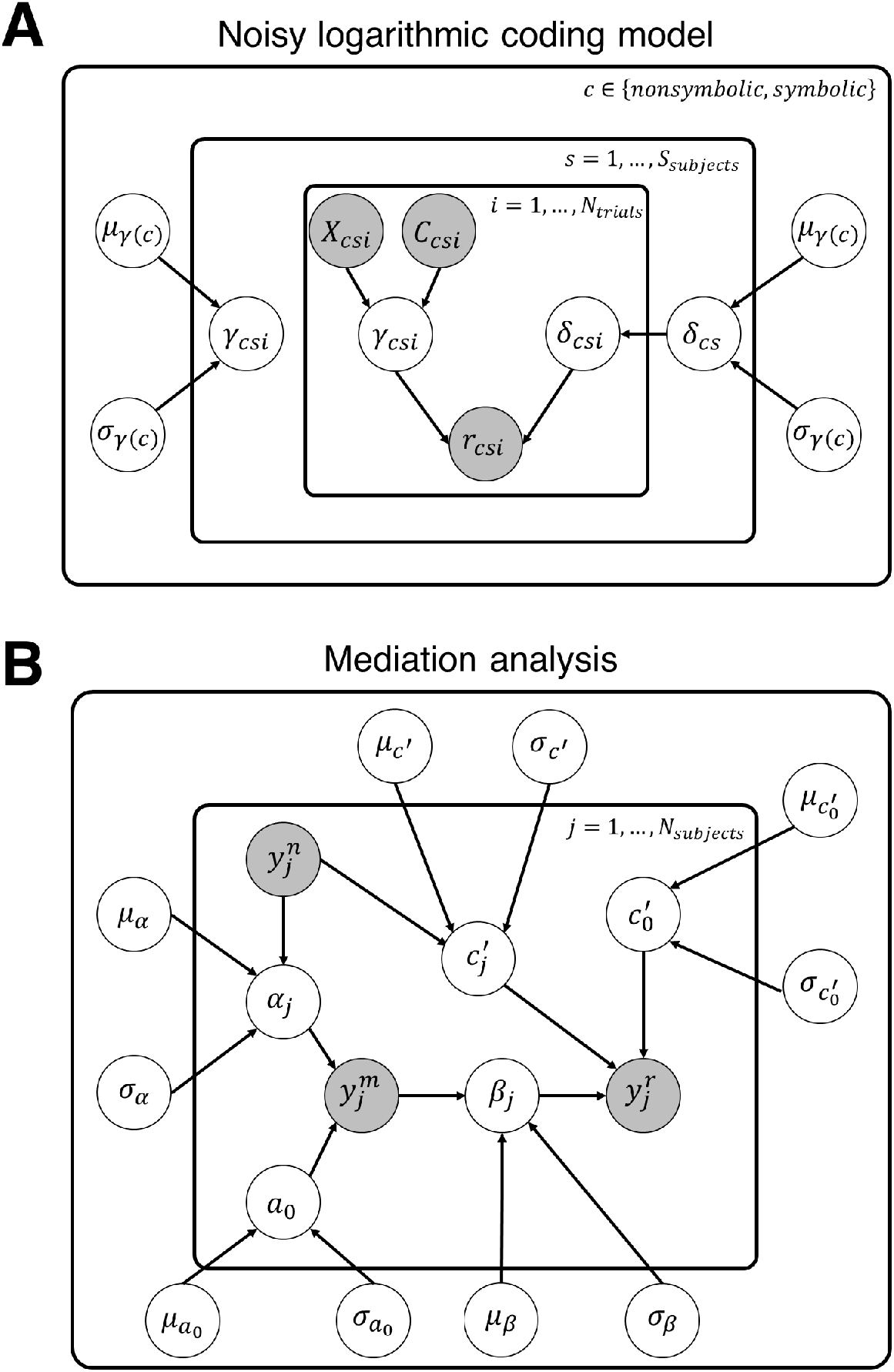
Hierarchical Bayesian models. Graphical representations of the hierarchical Bayesian **(a)** noisy logarithmic coding model and **(b)** mediation analysis. Clear circles represent latent variables while filled circles are observed variables, such as trialwise choice (*r*_*csi*_) data, subject-wise behavioural and neural measurements (*y*_*j*_), and numerosity / payoff inputs (*X, C*). The following equations show the distributions assumed for each of the model latent variables:

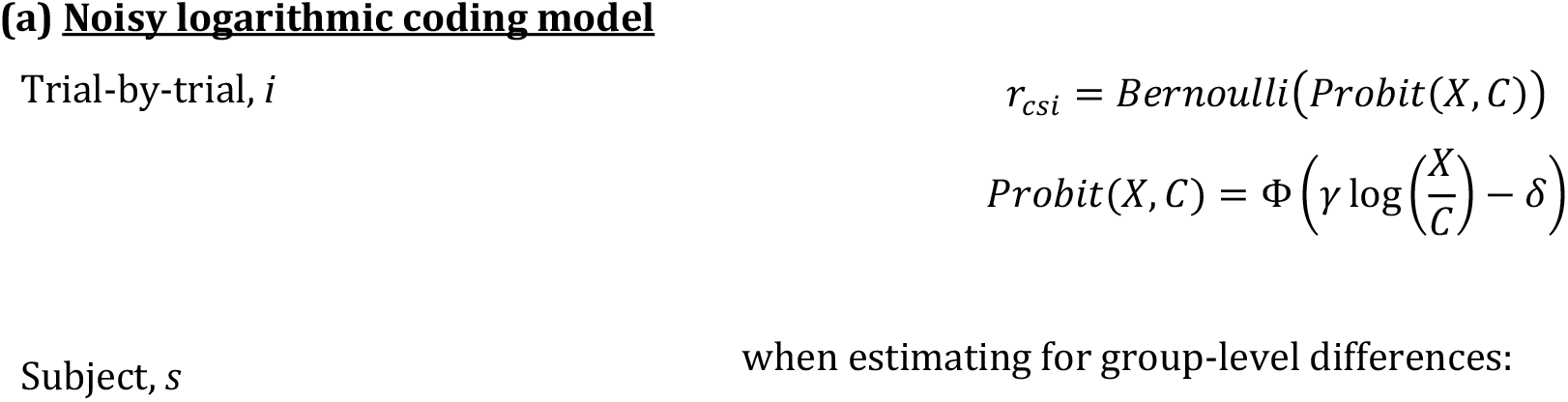
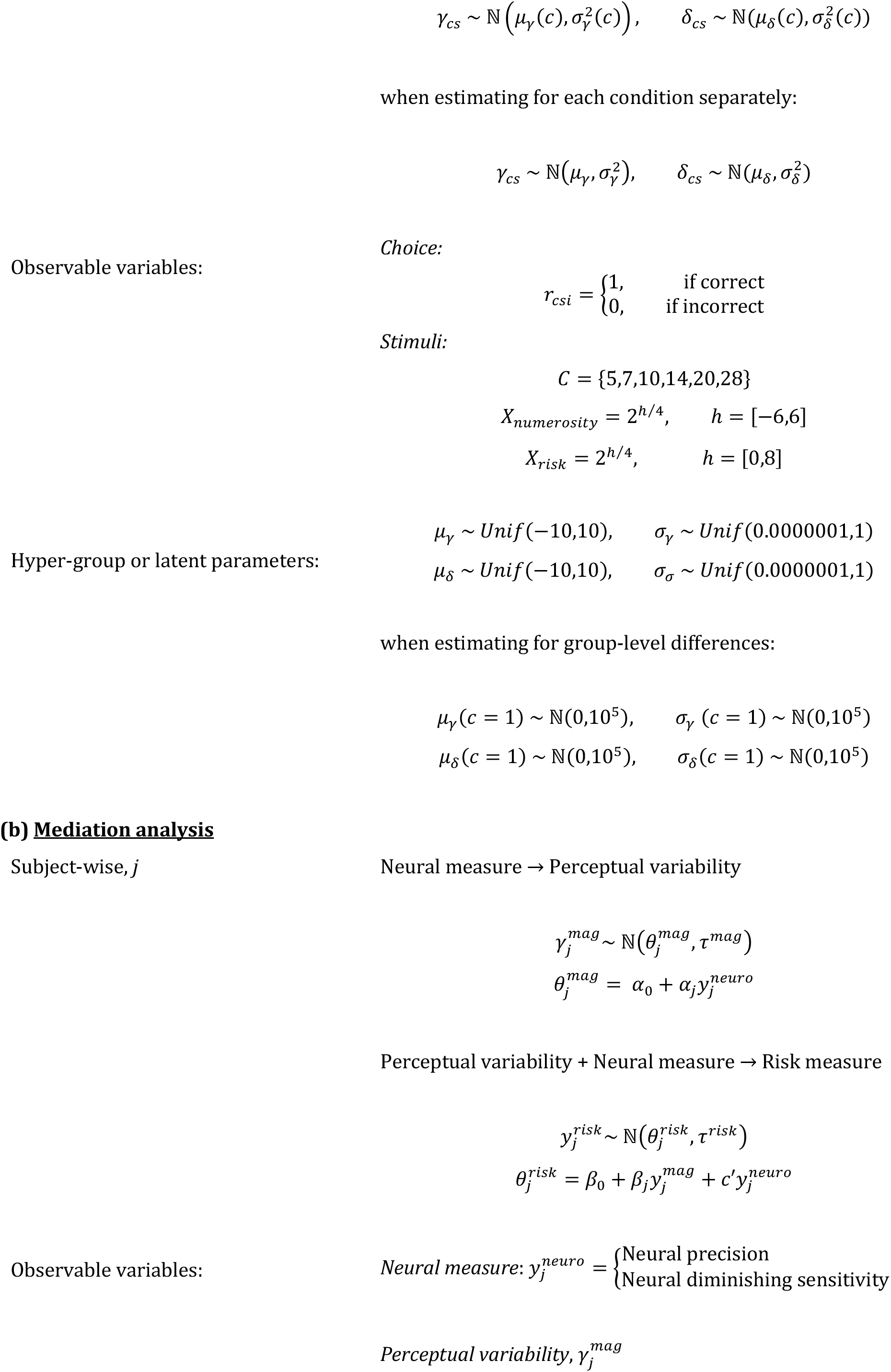
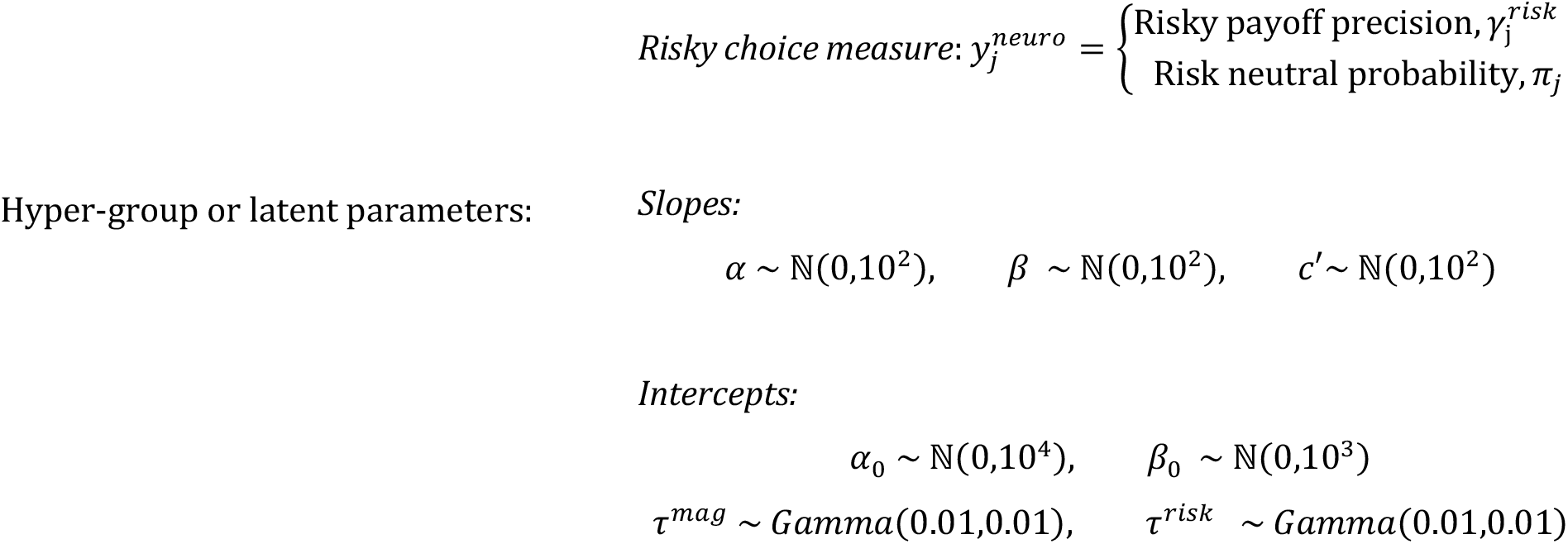

## Footnotes

1 The resulting Python package can be found on GitHub https://github.com/Gilles86/braincoder

